# Sequence-structured fitness landscapes predict influenza evolution

**DOI:** 10.64898/2026.10.27.762377

**Authors:** Edwin Rodríguez-Horta, John P. Barton

## Abstract

Seasonal influenza viruses continually evolve to evade host immunity. Vaccines must be regularly updated to protect against new circulating strains, motivating efforts to forecast viral evolution. Recent experimental and computational methods have made progress on this problem, but learning the fitness landscape driving viral evolution remains challenging because genomic surveillance samples only a small fraction of the possible sequence space. Here, we developed a generative framework that shares information across genetically related variants, inferring effective mutation-level contributions to viral fitness from temporal sequence data. Applied to hemagglutinin and neuraminidase from A/H3N2 and A/H1N1 viruses, the inferred fitness landscapes recovers known determinants of antigenicity and reveals convergent spatial organization across subtypes consistent with common antibody-mediated selection. Forward simulations from these landscapes predicted both the fixation or loss of circulating mutations and the future composition of viral populations, outperforming models based on neutral evolution or multinomial-logistic growth. Collectively, our results show that interpretable representations of pathogen fitness landscapes can reveal biologically important features and improve prospective evolutionary forecasting.

## Introduction

Seasonal influenza viruses cause substantial morbidity and mortality worldwide ^1,2^. Over time, these viruses accumulate mutations in the surface proteins hemagglutinin (HA) and neuraminidase (NA) that allow them to escape immunity generated by prior infection or vaccination ^3–6^. This continual antigenic drift necessitates frequent updates to influenza vaccines. Because vaccine strains must be selected months in advance – a constraint that can be relaxed, but not eliminated, with mRNA-based technology – these decisions depend on anticipating which viral variants will circulate in the future ^7,8^. Improving our ability to predict influenza evolution would therefore advance our understanding of viral adaptation and the rational selection of vaccine strains.

Efforts to predict influenza evolution have progressed from identifying positively selected sites and lineages to forecasting the future composition of circulating viral populations. Early studies used recurrent substitutions to identify lineages likely to persist ^9^, while subsequent models used fitness estimates based on antigenic change, mutational load, or phylogenetic branching patterns to predict successful clades and vaccine candidates ^10–12^. More recent approaches have integrated genetic and antigenic data, modeled site-specific mutation dynamics, or extrapolated lineage growth from temporal sequence data ^13–15^.

Here, we developed an interpretable, generative framework for forecasting influenza evolution using genomic surveillance data. Rather than assuming independent fitness values for viral lineages or clades, we represent variant fitness using shared contributions from their constituent amino acids. This sequence-structured model constrains similar viruses to generally have similar fitness, but it also allows specific mutations to drive substantial fitness differences when they are well-supported by data. Effective fitness parameters can be inferred from the patterns of viral variant frequency change over time ^16^. We then use the learned fitness model to forecast future evolution through stochastic simulations of infection transmission.

Applied to HA and NA from subtype H3N2 and H1N1 influenza A viruses (IAV), our framework readily identified positive selection in known antigenic and functional regions. We observed spatially convergent evolution across IAV subtypes in solvent-accessible regions, which may be driven by common antibody-mediated selection. In simulations, our model successfully predicted the fixation or loss of individual mutations, a task that was previously shown to be challenging even for strong phylogenetic predictors ^17^. Model forecasts of the future composition of the viral population also out-performed both simple neutral baseline predictions and ones generated by multinomial logistic regression (MLR), a state-of-the-art approach for viral fitness estimation and evolutionary prediction ^18,19^. While the present work focuses on IAV, our framework could be applied more broadly to interpret and predict evolution in other contexts.

## Results

### Modeling influenza virus transmission

We model influenza transmission as a branching process in which the effective reproduction number of each viral variant depends on its amino acid sequence (**Fig. 1**, Methods). Let variant *a* have sequence 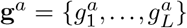 over a *q* = 20-state amino acid alphabet. Its effective reproduction number is *R*_*a*_ = *R* (1+*w*_*a*_), where *w*_*a*_ quantifies the relative fitness of variant *a*. Because only a small fraction of possible influenza sequences have been observed, the variant fitness model must generalize across nearby variants. However, it must also retain the ability to distinguish related variants with substantial differences in fitness. We thus represent variant fitness as

**Fig. 1.**
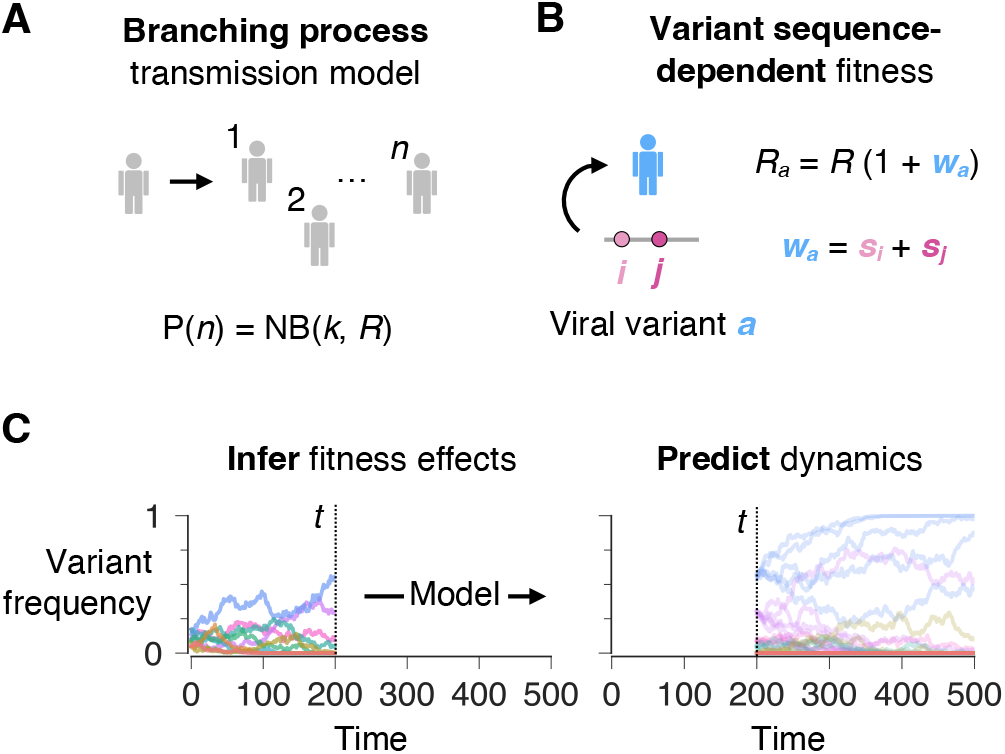
A sequence-structured generative framework for forecasting influenza evolution. (**A**) Viral transmission is modeled as a branching process. (**B**) The effective reproduction number *R*_*a*_ depends on the viral variant *a*, with additive contributions from each mutation. (**C**) To predict future evolutionary dynamics, we first estimate the distribution of likely fitness values for each variant from data collected until time *t*, parameterized by fitness effects for individual mutations (1). Then, we perform stochastic simulations with fitness sampled from the inferred distribution, starting from the state of the population at time *t*.

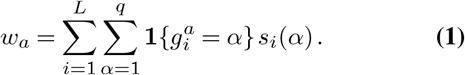

The resulting parameters *s*_*i*_(*α*) can be interpreted as effective selection coefficients. This representation shares statistical strength across related sequences: genetically similar variants will typically have similar fitness unless evidence suggests that the mutations that separate them have large fitness effects. In principle, the *w*_*a*_ and *s*_*i*_(*α*) could vary with time, for example reflecting changes in population immunity. While this case remains computationally tractable ^20^, here we focus on the biological insights and predictions that can be generated from a constant fitness model.

Directly fitting the branching process model to data is challenging, so we consider a diffusion limit analogous to that in population genetics ^21–23^. We assume that the number of infected individuals *N* is large and the fitness differences between variants are small (|*w*_*a*_|*≪* 1). In this limit, the likelihood of an evolutionary history or “path,” described by the distribution of variant frequencies over time, is a simple function of the fitness parameters *s*_*i*_(*α*). With a zero-mean Gaussian prior of precision *γ*, the maximum a posteriori (MAP) estimate of mutation fitness contributions is ^16^

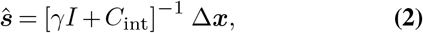

where Δ***x*** is the vector of net changes in mutant frequencies over the observation window and *C*_int_ is the time-integrated covariance of mutant allele frequencies. Here, other epidemiological parameters such as the population size *N* have been effectively absorbed into the regularization strength *γ* ^16^. Similar expressions for estimating selection have also been derived from population genetics models ^23^.

Equation (2) combines two complementary sources of information. The term Δ***x*** identifies mutations that increase or decrease in frequency over the observation period. However, mutations do not evolve independently. The covariance matrix *C*_int_ captures the co-occurrence of mutations across circulating variants, allowing the model to separate the effects of mutations from the genetic background. Regularization controls spurious inferences by shrinking weakly constrained mutation fitness contributions toward zero.

### Simulations to forecast evolution

The branching process described above is a *generative* model for viral transmission at the population level. Thus, we can use estimated fitness effects *s*_*i*_(*ω*) and knowledge of current variant frequencies *x*_*a*_ to simulate changes in the viral population across infected individuals over time. We incorporate uncertainty into our predictions by performing multiple simulations using fitness effects sampled from the posterior distribution, centered on (2) (Methods).

We use these simulations to generate two complementary forecasts. First, we estimate the relative probability that each circulating mutation ultimately fixes in the population. Second, we predict the composition of the viral population at a specified future time by selecting the medoid of *M* = 100 simulated populations. The medoid is defined as the population that has the smallest total distance to all other simulated populations, thus making this population a good representative of typical future evolution. Here, we used earth mover’s distance (EMD) as a distance metric (Methods).

By incorporating both individual mutation effects and population variation, our simulations capture the importance of genetic context in evolution. For example, beneficial mutations that arise on a weak genetic background may be lost as that variant is outcompeted by others that are more transmissible. Conversely, neutral mutations can hitchhike to fixation on an advantageous background.

### Validation in simulations

We first validated our approach on synthetic data generated by the branching process model. Our simulations evolved a population of around *N* = 1000 infected individuals over 1000 rounds of transmission with randomly generated fitness effects (Methods). Across replicate simulations, inferred selection coefficients closely matched their true values (**Fig. S1A**).

To benchmark our predictive power, we compared model forecasts to those from a simple, neutral model. In neutral evolution, where mutations have no effects on fitness, the probability of a mutation fixing is simply equal to its current frequency. The neutral prediction for the future distribution of variant frequencies in the viral population is the same as the current distribution at the time that the prediction is made.

In simulations, our model substantially exceeded the accuracy of the neutral model in predicting the fates of individual mutations, especially ones at intermediate frequencies (**Fig. S1B-C**). As expected, predictions of the future composition of the viral population declined as the time horizon for forecasting increased. However, model predictions remained closer to the true distribution than the neutral model for all the time horizons that we tested (**Fig. S1D**).

### Selection at functional sites in IAV surface proteins

Having validated our approach in simulations, we next asked whether our model could recover known biological determinants of influenza evolution. We analyzed amino acid sequences for HA and NA from IAV subtypes A/H3N2 and A/H1N1pdm deposited in GISAID ^27^. We focused on sequences collected between early 2009 and March, 2020, over which time IAV sequences were densely sampled. After quality control filtering, our final data set contained roughly 50,000 sequences for each protein and IAV subtype (**Fig. S2**, Methods). Multiple clusters of mutations rise and fall during this time (**Fig. S3A**).

The inferred distribution of mutation fitness contributions was strikingly similar across all IAV surface proteins and subtypes (**Fig. S3B**). Most mutations clustered near neutrality, consistent with either small fitness effects or limited information to infer strong selection. We observed a long tail of rare beneficial mutations, reaching a maximum selection coefficient of around 1% per day. In contrast, very few mutations were inferred to be strongly deleterious.

If the inferred fitness landscape captures biologically meaningful selection, beneficial mutations should be enriched at known antigenic and functional sites. As one example, the HA receptor binding site (RBS) is critical for cell entry, and mutations in or near RBS can also drive changes in antigenicity ^5,6,26^. Consistent with our expectations, we found that mutations within both RBS and well-characterized antigenic sites were significantly more beneficial than mutations elsewhere in the protein (**Fig. 2A-B**).

**Fig. 2.**
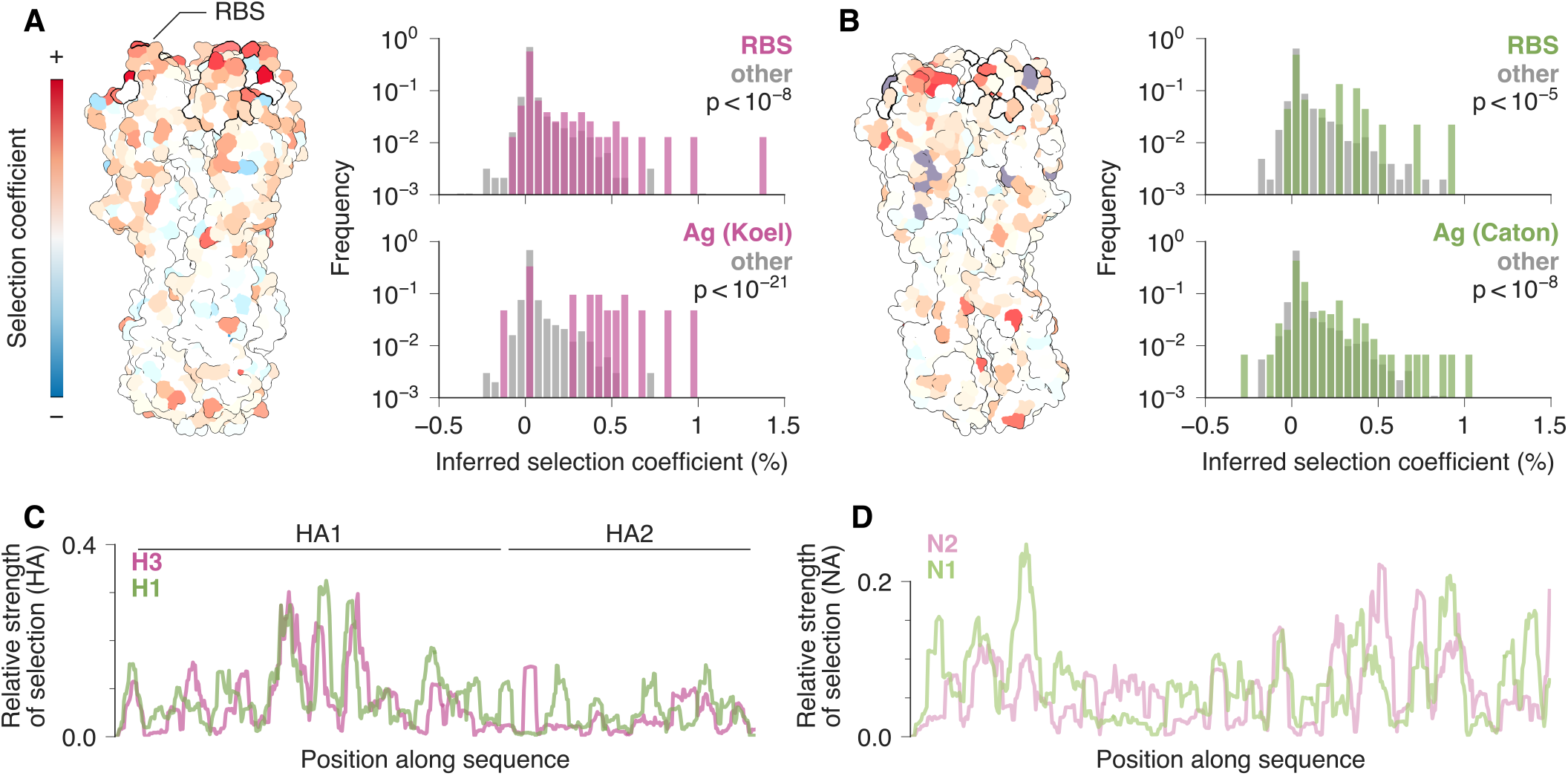
The inferred fitness landscape recovers functional regions and reveals convergent selection across influenza subtypes. (**A**) Inferred mutation fitness contributions for H3 mutations mapped onto the protein structure ^24^ (PDB: 4WE8). Mutations in the receptor binding site (RBS) and identified antigenic sites ^5^ are more strongly selected than mutations in other regions (Student’s t-test). (**B**) Map of fitness contributions for H1 mutations on the protein structure ^25^ (PDB: 3LZG). Like H3, H1 mutations in RBS and antigenic sites ^26^ were inferred to be more beneficial than those at other sites, on average. (**C**) In HA1, we find similar patterns of selection across H3 and H1 that also strongly correlate with solvent accessible surface area, likely due to antibody exposure. Patterns of selection in HA2 and (**D**) NA are more idiosyncratic.

The strongest individual fitness effects also typically corresponded to mutations with experimentally characterized phenotypes (**Tables 1-4**). In H3, strongly selected mutations clustered in antigenic sites surrounding the receptor-binding site (e.g., T131K, N189K, and F159Y), including residues 159 and 189, which are major determinants of H3N2 antigenic evolution ^5,28^. In H1, K166Q, one of our three most strongly selected HA mutations, is a key antigenic substitution associated with reduced recognition by human antibodies during the 2013–2014 H1N1 season ^29^, while N129D has been shown to confer antibody escape and increase viral replication in vitro ^30^. Our most strongly selected mutation in N2, E344K, reduces inhibition by antisera against earlier H3N2 viruses ^31^. Strikingly, several of the strongest N2 and N1 mutations cluster near the second sialic acid-binding site (2SBS): N369K and K432E in N1 have experimentally demonstrated effects on both NA antigenicity and receptor processing activity, with K432E driving a particularly large antigenic change ^32^. Mutations in 2SBS have previously been identified as contributors to inter-gene epistasis, potentially mediating HA-NA balance ^33,34^. Together, these examples reveal a consistent pattern in which the largest fitness contributions lie in regions that mediate antibody recognition, receptor binding, or NA–receptor interactions.

### Convergence of selection across IAV subtypes

Having inferred fitness landscapes independently for each subtype, we next asked whether these landscapes shared common spatial organization. Such convergence would suggest that conserved structural or functional constraints shape IAV adaptation. To this end, we compared smoothed maps of the relative strength of selection along each protein sequence (**Fig. 2C-D**, Methods). Despite substantial sequence divergence between IAV subtypes, we observed a strong alignment between H3 and H1 landscapes (Pearson’s *r* = 0.58, *p <* 10^*−*50^), which was primarily driven by correlations within HA1 (Pearson’s *r* = 0.62, *p<* 10^*−*36^). In contrast, little correlation was observed for HA2, the stalk region (Pearson’s *r* = − 0.02). N2 and N1 exhibited similar convergence, but with a more modest correlation compared to similarities across subtypes in HA (Pearson’s *r* = 0.28, *p<* 10^*−*9^).

The strong correspondence between H3 and H1 within HA1 suggested that selection for antibody escape may drive common spatial patterns across subtypes. Because antibody epitopes are concentrated on exposed protein surfaces, we reasoned that selection should be correlated with solvent accessibility. Indeed, solvent accessible surface area (SASA, smoothed in the same way along the protein sequence; Methods) was strongly associated with selection in HA, especially in HA1 (Pearson’s *r* = 0.64 and 0.46, *p<* 10^*−*37^ and 10^*−*17^ for H3 and H1, respectively; **Fig. S4**). Correlations between selection and SASA in NA, where structural data was available, were similar to those in HA (Pearson’s *r* = 0.59 and 0.37, *p<* 10^*−*37^ and 10^*−*13^ for N2 and N1, respectively). The association between SASA and selection was stronger for H3N2 than for H1N1, consistent with its higher rate of antigenic evolution ^35^. These results demonstrate a common spatial organization of positive selection across IAV subtypes, consistent with antibody-mediated immune pressure in exposed regions of viral surface proteins.

### Forecasting the fixation or loss of mutations

Above, we characterized the inferred IAV fitness landscape within and across virus subtypes. Next, we tested its ability to predict future evolutionary dynamics. Every 10 days, starting from 3 years after the initial time point, we inferred the fitness landscape using only sequences collected before the prediction date and simulated future viral evolution (**Fig. 1C**, Methods). No information from subsequent time points was used during inference or forecasting.

We first assessed the ability of the model to predict the evolutionary fate of circulating mutations. In **Figure 3**A, we group mutations into bins based on their frequency in the global IAV population at the prediction time and stratify them according to their relative likelihood of fixation in our simulations. In a neutral model, the probability of fixation for a mutation is simply equal to its frequency. The average fixation probability across all mutations is close to this expectation, consistent with previous observations that influenza evolution appears nearly neutral at the level of individual mutations ^17^.

**Fig. 3.**
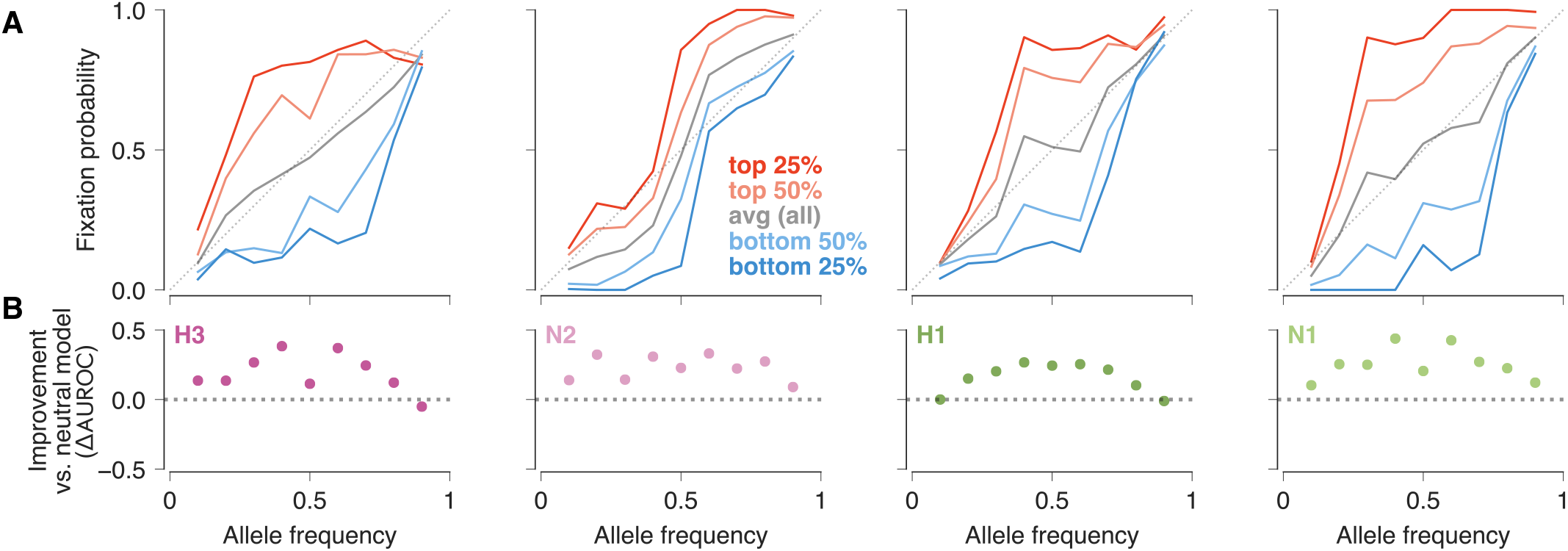
The inferred fitness landscape predicts the evolutionary fate of circulating mutations. (**A**) Mutation frequency versus the fraction of mutations that ultimately fix in the viral population (cf. ^17^). Mutations are stratified according to their relative likelihood of fixation in simulations, from those predicted to be the most likely to fix (top 25%) to the least (bottom 25%). (**B**) Improvement in AUROC relative to the neutral model. The relative predictive power of our model is typically highest at intermediate frequencies. Full AUROC scores are shown in **Fig. S5.**

However, our model successfully discriminates between mutations that are likely to fix and those that are likely to be lost. We quantified prediction accuracy using the area under the receiver operating characteristic (AUROC; Methods) for each frequency bin (**Fig. 3B, Fig. S5**). We found the greatest predictive performance for mutations at intermediate frequencies. Predictions for mutations at very low or high frequencies were closer to those obtained from a simple, neutral model. Importantly, our simulations preserve the genotypes carrying each mutation, allowing the model to use both information about the focal mutation and its genetic background to predict fixation or loss.

### Forecasting future viral populations

Beyond the fates of individual mutations, we also generated forecasts for the future sequence composition of the viral population (**Fig. S6**, Methods). This challenging prediction problem could be particularly informative for vaccine design ^13^, with the goal of matching vaccine strains to those that are likely to be common in the next season. To assess our predictive power, we compared model predictions to the true distribution of variant frequencies in the viral population using earth mover’s distance (EMD) ^13,36^, an average Hamming distance between sequence distributions weighted by variant frequencies (Methods). We considered a range of prediction time horizons, forecasting the composition of the viral population 6, 9, and 12 months into the future.

Across both IAV subtypes and surface proteins, our simulated forecasts were substantially closer to the true future population than neutral predictions (**Figure 4**). Improvements were especially consistent for H1N1. Similar results were observed for all prediction horizons, with a slight tendency toward greater performance improvements at longer horizons (**Figs. S7-S8**).

**Fig. 4.**
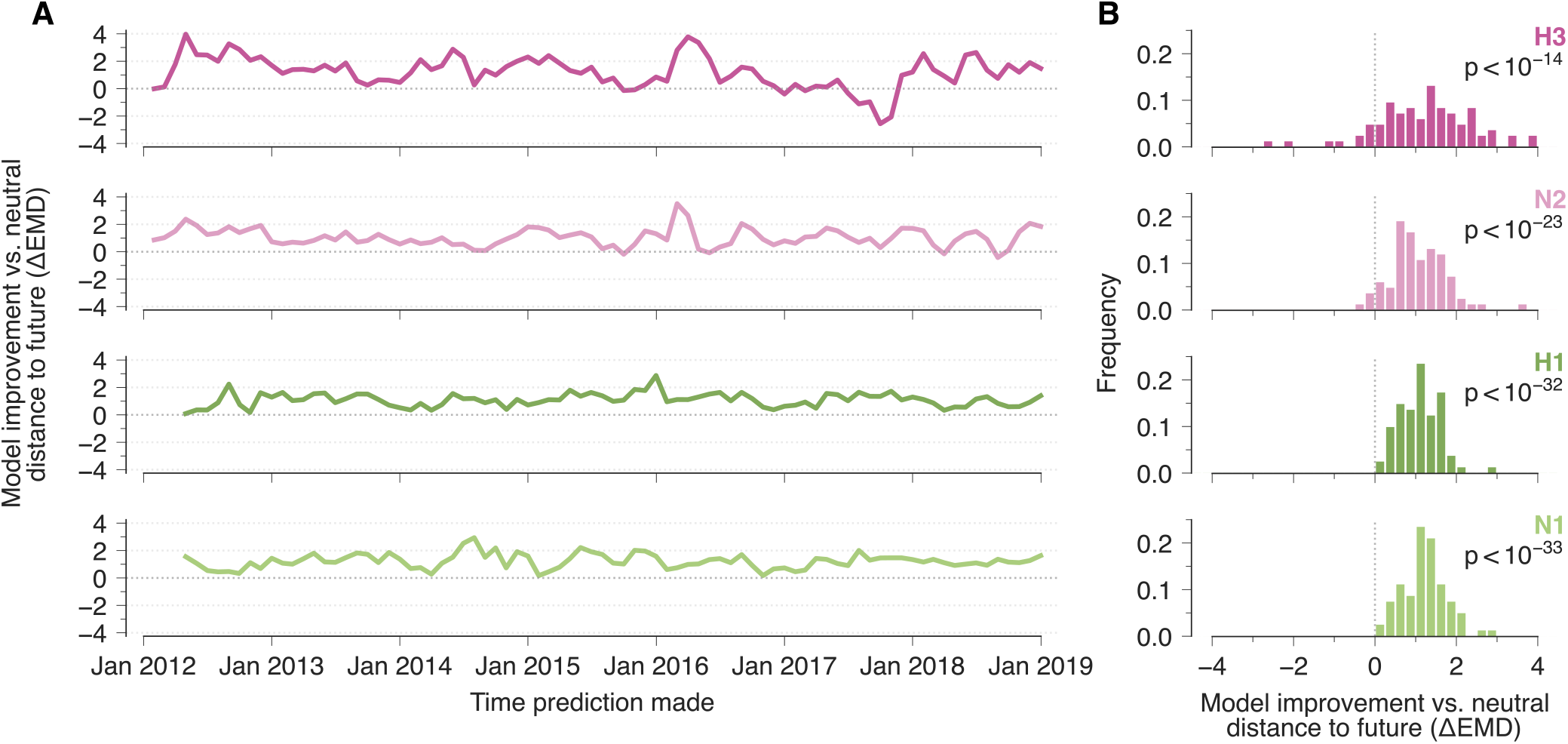
Sequence-structured forecasts accurately predict future influenza populations. (**A**) Our simulations yield more accurate predictions of IAV sequence composition 9 months in the future than a neutral model, as measured by the difference in earth mover’s distances (ΔEMD) between the true population and neutral vs. simulated ones. **(B)**Histogram of ΔEMD values. For each IAV subtype and surface protein, our simulations improve significantly over neutral ones (Student’s t-test). Comparison against the neutral model at 6 and 12 month time horizons are shown in **Figs. S7-S8**, and comparisons against MLR are shown at 6, 9, and 12 month time horizons in **Figs. S9-S11**.

We next compared our forecasts with multinomial logistic regression (MLR; Methods), a widely used approach for estimating viral fitness and forecasting evolution ^18,19^. Our simulations provided significantly more accurate forecasts of the future IAV population, on average, for every IAV subtype, surface protein, and time horizon that we tested (**Figs. S9-S11**). These results demonstrate the strength of our approach in forecasting near-future IAV evolution.

Notably, however, there were instances where our model underperformed. In particular, evolutionary forecasts for H3 made in mid-2017 were less accurate than neutral ones. The 2017-2018 influenza season was characterized by a high disease burden, despite the dominance of 3C.2a lineage H3N2 viruses in both 2016-2017 and 2017-2018^37^. Although 3C.2a2 lineage viruses were declining in early to mid 2017, reassortment yielded a new variant with the 3C.2a2 HA and PB1 gene segments paired with other gene segments from 3C.2a1 viruses, which expanded dramatically in early 2018^38^. Experimental studies showed that the reassortant virus replicated more efficiently than parental 3C.2a1 and 3C.2a2 viruses and was associated with more severe infections ^38^. Because this increase in fitness arose from reassortment involving genome segments *outside* HA, it could not be anticipated from HA sequence evolution alone. This example illustrates an important limitation of sequence-based forecasting restricted to a single viral gene segment.

### A sequence-structured variant fitness model improves evolutionary forecasts

The success of a circulating mutation may depend on both its own fitness contribution and the sequence background on which it occurs. To determine how these factors contribute to evolutionary predictability, we compared mutations that ultimately fixed with those that were lost. Overall, mutations that fixed typically had larger inferred fitness contributions (**Fig. 5A, Fig. S12**, Methods). However, their success could not be explained by the focal mutation alone. Mutations that fixed tended to appear on fitter backgrounds compared to the population average, and the fitness of those backgrounds subsequently increased more rapidly than for backgrounds carrying mutations that were lost (**Fig. 5B-C, Figs. S13-14**).

**Fig. 5.**
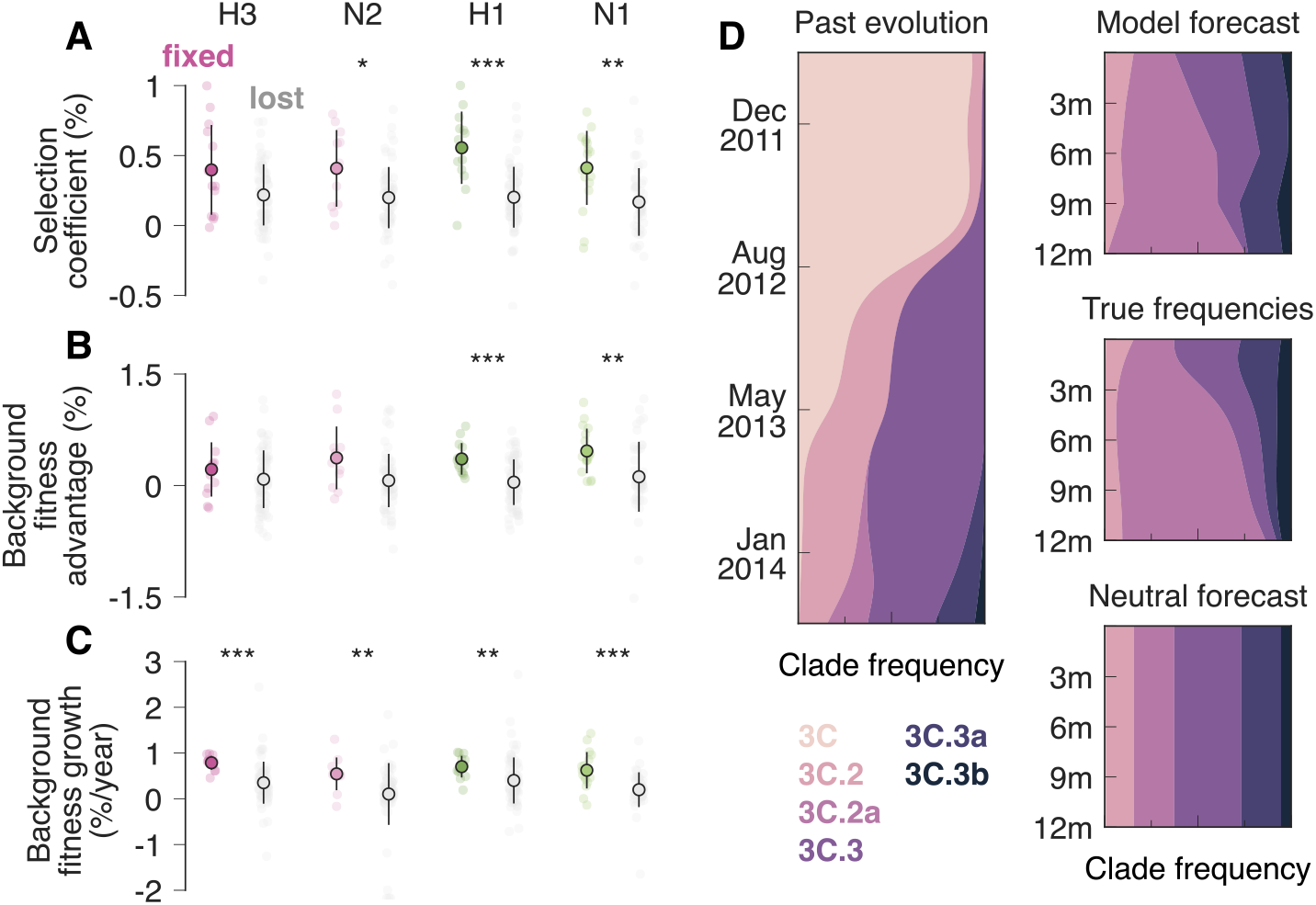
Mutation effects and linked sequence backgrounds jointly shape influenza evolution. (**A**)Mutations that fix tended to have larger fitness contributions than mutations that were lost, although the strength of this association varied among proteins. (**B**) Mutations that fixed also generally appeared on fitter backgrounds, as measured by the average fitness of sequences carrying the focal mutation relative to the population average when the mutation first crossed 5% frequency. (**C**) Backgrounds carrying mutations that fixed tended to gain fitness more rapidly. *: *p<* 0.05, **: *p<* 0.01, ***: *p<* 0.001, Student’s t-test. (**D**) Example of predicted H3 clade dynamics. The model was trained on the sequence population observed through June 2014 and used to forecast population composition 3, 6, 9, and 12 months into the future. We grouped sequence-level forecasts into established phylogenetic clades after simulation for visualization. The model captured the expansion and contraction of major clades observed in the future population more closely than a neutral forecast that preserved their frequencies at the prediction date.

We thus hypothesized that discarding genetic background information should reduce predictive power. To test this hypothesis, we applied our approach to single mutations alone, ignoring the background sequence when inferring fitness effects and simulating future evolution (Methods). Consistent with our expectations, while this simple model still surpassed neutral fixation/loss predictions, it performed worse than our full model (**Fig. S15**). Retaining complete protein sequences also consistently improved forecasts of population-level H3 evolution (**Fig. S16-18**). Interestingly, however, the single site model yielded slightly better predictions for H1 and N1 at longer time horizons, demonstrating that a simple model could also generate effective predictions of future sequence composition.

Conversely, increasing the flexibility of the fitness model did not necessarily improve predictions. Typically, models similar to MLR ^18,19^ assign separate growth parameters to circulating lineages or variants, as implemented here. As a result, nearby sequences need not have similar fitness estimates, and sparsely sampled lineages cannot borrow information from related sequences. In contrast, our fitness model (1) shares information across related variants while allowing small numbers of mutations to drive substantial fitness differences when well-supported by data. For the forecasting tasks we considered, this intermediate representation outperformed the more flexible variant-level one across IAV subtypes, proteins, and time horizons (**Figs. S9-S11**).

To illustrate how sequence-level forecasts translate into familiar patterns of influenza evolution, we grouped predicted H3 variants into established phylogenetic clades after simulation (**Fig. 5D**). Clade labels were used only for visualization and played no role in model fitting or forecasting. Starting from the viral population observed in June 2014, the model reproduced the subsequent expansion of 3C.2a and the contraction of competing lineages, while the neutral forecast retains the clade composition present at the prediction date. Thus, large-scale patterns of clade turnover emerged from the sequence-level fitness landscape and competition among sequence backgrounds, without requiring clades to be defined as the units of prediction.

## Discussion

Here, we developed a generative framework for learning from and forecasting influenza evolution using genomic surveillance data. In applications to HA and NA sequences from A/H3N2 and A/H1N1 viruses, the inferred fitness landscapes recovered strong selection in known antigenic and functional regions. Our results also revealed convergent spatial organization across subtypes, particularly at solvent-exposed sites in the HA head. At the mutation level, several of the mutations with the largest inferred effects displayed experimentally characterized changes in antibody recognition, glycan shielding, receptor avidity, or NA receptor processing, including a notable concentration of selected N1 and N2 substitutions on the 2SBS surface. Stochastic simulations of viral evolution predicted both the fixation or loss of circulating mutations and the future composition of viral sequence populations. Our model showed significant improvements in predictive power compared to both a simple baseline (neutral evolution) and more flexible multinomial logistic regression. A simplified single site version of our approach also displayed impressive predictive performance, occasionally out-performing the full model. Collectively, these results show that an interpretable approximation to the influenza fitness landscape can capture drivers of past adaptation and forecast future evolution.

Our framework builds on a long history of influenza forecasting and fitness inference. Previous approaches have predicted successful viral lineages using genetic, phylogenetic, and phenotypic features ^9–13,39^. More recent methods have modeled the dynamics of competing residues at individual sites ^14^ or coupled adaptive sequence evolution to birth–death phylodynamics ^40^. Our approach combines several complementary features: it infers effective fitness from temporal distributions of protein sequences ^16,23^, operates in sequence space without requiring predefined clades, and shares information across related variants through a structured representation of fitness. Forward simulations also incorporate both parameter uncertainty and evolutionary stochasticity.

For influenza forecasts based on genomic surveillance, it can be challenging to learn a mapping from viral sequence to fitness from data that sample only a small fraction of the possible sequence space. Representations based on discrete viral lineages or clades impose hard boundaries, assigning identical fitness within groups with potentially unconstrained differences between them. Our fitness representation instead imposes a soft constraint, such that genetically similar viruses are expected, but not required, to have similar fitness. Conceptually, this model should be viewed as a controlled approximation rather than a mechanistic description of influenza fitness. The fitness effects that we infer depend on the genetic and epidemiological contexts present in the data, and the model assumes constant, additive fitness contributions despite known effects of changing immune environments and epistasis ^3,41,42^. Although richer models could represent these processes explicitly, additional biological complexity may not improve prediction when data is limiting. Our results suggest that effective forecasting can arise from representations that capture the dominant, learnable structure of fitness across a sparsely sampled sequence space.

Genetic background has long been recognized to play an important role in influenza evolution. Experimental studies identified mutations that enabled the emergence of oseltamivir resistance ^41^, demonstrated stability-mediated epistasis ^42^, and revealed coordinated evolution between HA and NA ^33^, while population genetic analyses showed that linkage and clonal interference strongly influence mutation fate ^43,44^. Consistent with these observations, we found that mutations that fixed tended to occur on fitter sequence backgrounds whose fitness continued to increase during subsequent evolution. Even without the explicit inclusion of epistasis in our model, competition between variants allows background fitness to affect evolutionary trajectories.

Despite substantial sequence divergence between H3N2 and H1N1, they shared similar spatial patterns of selection. Previous studies have shown that adaptive evolution is concentrated in the globular head of HA, particularly at solvent-exposed antigenic regions near the receptor binding site ^3,45,46^, and that these regions are unusually tolerant to mutations ^47^. We found that this organization extends more broadly across subtypes, and with a slightly weaker but significant correspondence in NA. However, we note that similar spatial patterns in selection do not necessarily reflect convergent fitness at the amino acid level. HA subtypes can maintain similar structures despite substantially different sequence constraints and amino acid preferences ^48^, and antigenic evolution is constrained by pleiotropic effects on stability and function ^49^. Rather, the association between selection and solvent accessibility that we observe suggests that subtype-specific constraints could be superimposed with common pressure from antibody exposure.

Comparison of strongly selected mutations further sharpened these spatial patterns. In H3, several highly selected substitutions have been linked to altered antigenicity or receptor avidity, including N145S, F193S, and T135K (refs. ^28,50^). Correspondingly in H1, N129D and S183P have direct effects on antibody recognition or receptor binding ^30,51^; the adjacent S162N, K163Q, and S164T substitutions also trace remodeling of the immunodominant N162 glycosylation site ^52^. In NA, several of the most strongly selected substitutions clustered on the second sialic acid-binding site surface: N2 S367N, K369T, and N402D occupy the 370- and 400-loop regions, and N1 N369K and K432E experimentally affect both receptor processing and antigenicity ^32^; N386K partly compensates the effect of K432E. Overall, this concentration is consistent with selection on both immune recognition and HA–NA–receptor balance ^33,34^.

The framework that we have described also has limitations. While our current analyses treat each influenza surface protein independently, the 2017 H3N2 reassortment event ^38^ shows that lineage success can depend on genetic variation elsewhere in the genome. Reassortment and coordinated evolution among influenza gene segments have been observed in multiple instances ^33,53,54^, motivating an extension to model whole genomes. Our current simulations also consider only circulating viral variation at the prediction time. Incorporating novel mutations and reassortment could help to more realistically model uncertainty, especially for longer forecasts. Finally, evolutionary forecasting addresses only one component of vaccine strain selection, which is also informed by antigenic characterization, manufacturing constraints, and other public health considerations.

Expanding pathogen genomic surveillance creates new opportunities to understand and anticipate evolution. Our results show that an approximate representation of the fitness landscape can be inferred from such data and used to successfully forecast future evolution. Extending similar approaches across whole genomes, changing selective environments, and other rapidly evolving pathogens may further improve our ability to quantitatively understand pathogen evolution.

## ACKNOWLEDGEMENTS

The work of E.R-H. and J.P.B. reported in this publication was supported by the National Institute of General Medical Sciences of the National Institutes of Health under Award Number R35GM161641. The content is solely the responsibility of the authors and does not necessarily represent the official views of the National Institutes of Health.

## AUTHOR CONTRIBUTIONS

All authors designed research, performed research, analyzed data, contributed new reagents/analytic tools, and wrote the paper.

## References

1. Iuliano, A. D. et al. Estimates of global seasonal influenza-associated respiratory mortality: a modelling study. The Lancet 391, 1285–1300 (2018).

2. Organization, W. H. https://www.who.int/news-room/fact-sheets/detail/influenza-(seasonal) (Accessed Feb 16, 2023.). URL https://www.who.int/news-room/fact-sheets/detail/influenza-(seasonal).

3. Petrova, V. N. & Russell, C. A. The evolution of seasonal influenza viruses. Nat Rev Microbiol 16, 47–60 (2017).

4. Smith, D. J. et al. Mapping the antigenic and genetic evolution of influenza virus. science 305, 371–376 (2004).

5. Koel, B. F. et al. Substitutions near the receptor binding site determine major antigenic change during influenza virus evolution. Science 342, 976–979 (2013).

6. Wu, N. C. & Wilson, I. A. Influenza hemagglutinin structures and antibody recognition. Cold Spring Harbor perspectives in medicine 10, a038778 (2020).

7. Klingen, T. R., Reimering, S., Guzmán, C. A. & McHardy, A. C. In silico vaccine strain prediction for human influenza viruses. Trends Microbiol 26, 119–131 (2017).

8. Morris, D. H. et al. Predictive modeling of influenza shows the promise of applied evolutionary biology. Trends Microbiol 26, 102–118 (2017).

9. Bush, R. M., Bender, C. A., Subbarao, K., Cox, N. J. & Fitch, W. M. Predicting the evolution of human influenza a. Science 286, 1921– 1925 (1999). URL https://www.science.org/doi/abs/10.1126/science.286.5446.1921. https://www.science.org/doi/pdf/10.1126/science.286.5446.1921.

10. Fluksza, M. & Lässig, M. A predictive fitness model for influenza. Nature 507, 57 (2014).

11. Neher, R. A., Russell, C. A. & Shraiman, B. I. Predicting evolution from the shape of genealogical trees. eLife 3, e03568 (2014). URL 10.7554/eLife.03568.

12. Steinbrück, L., Klingen, T. R. & McHardy, A. C. Computational prediction of vaccine strains for human influenza a (H3N2) viruses. J Virol 88, 12123– 12132 (2014).

13. Huddleston, J. et al. Integrating genotypes and phenotypes improves longterm forecasts of seasonal influenza a/h3n2 evolution. eLife 9, e60067 (2020). URL 10.7554/eLife.60067.

14. Lou, J. et al. Predictive evolutionary modelling for influenza virus by site-based dynamics of mutations. Nature Communications 15, 2546 (2024).

15. Du, X., King, A. A., Woods, R. J. & Pascual, M. Evolution-informed forecasting of seasonal influenza a (h3n2). Science translational medicine 9, eaan5325 (2017).

16. Lee, B. et al. Inferring effects of mutations on sars-cov-2 transmission from genomic surveillance data. Nature Communications 16, 441 (2025).

17. Barrat-Charlaix, P., Huddleston, J., Bedford, T. & Neher, R. A. Limited predictability of amino acid substitutions in seasonal influenza viruses. Mol Biol Evol 38, 2767–2777 (2021).

18. Annavajhala, M. K. et al. Emergence and expansion of sars-cov-2 b. 1.526 after identification in new york. Nature 597, 703–708 (2021).

19. Abousamra, E., Figgins, M. & Bedford, T. Fitness models provide accurate short-term forecasts of sars-cov-2 variant frequency. PLOS Computational Biology 20, e1012443 (2024).

20. Gao, Y., Lee, B. & Barton, J. P. Inferring fitness seascapes from evolutionary histories. bioRxiv (2025).

21. Ewens, W. J. Mathematical Population Genetics 1: Theoretical Introduction (Springer Science & Business Media, 2012).

22. Mustonen, V. & Lässig, M. Fitness flux and ubiquity of adaptive evolution. Proceedings of the National Academy of Sciences 107, 4248–4253 (2010).

23. Sohail, M. S., Louie, R. H. Y., McKay, M. R. & Barton, J. P. MPL resolves genetic linkage in fitness inference from complex evolutionary histories. Nat Biotechnol 39, 472–479 (2020).

24. Yang, H. et al. Structure and receptor binding preferences of recombinant human a (h3n2) virus hemagglutinins. Virology 477, 18–31 (2015).

25. Xu, R. et al. Structural basis of preexisting immunity to the 2009 h1n1 pandemic influenza virus. Science 328, 357–360 (2010).

26. Caton, A. J., Brownlee, G. G., Yewdell, J. W. & Gerhard, W. The antigenic structure of the influenza virus a/pr/8/34 hemagglutinin (h1 subtype). Cell 31, 417–427 (1982).

27. Shu, Y. & McCauley, J. GISAID: Global initiative on sharing all influenza data from vision to reality. Euro Surveill 22 (2017).

28. Chambers, B. S., Parkhouse, K., Ross, T. M., Alby, K. & Hensley, S. E. Identification of hemagglutinin residues responsible for h3n2 antigenic drift during the 2014–2015 influenza season. Cell Reports 12, 1–6 (2015).

29. Linderman, S. L. et al. Potential antigenic explanation for atypical h1n1 infections among middle-aged adults during the 2013–2014 influenza season. Proceedings of the National Academy of Sciences 111, 15798–15803 (2014).

30. Yasuhara, A. et al. Diversity of antigenic mutants of influenza a (h1n1) pdm09 virus escaped from human monoclonal antibodies. Scientific reports 7, 17735 (2017).

31. Ge, J. et al. The antibody response against neuraminidase in human influenza a (h3n2) virus infections during 2018/2019 flu season: Focusing on the epitopes of 329-n-glycosylation and e344 in n2. Frontiers in Microbiology 13, 845088 (2022).

32. Dai, M. et al. Analysis of the evolution of pandemic influenza a (h1n1) virus neuraminidase reveals entanglement of different phenotypic characteristics. MBio 12, 10–1128 (2021).

33. Neverov, A. D., Kryazhimskiy, S., Plotkin, J. B. & Bazykin, G. A. Coordinated evolution of influenza a surface proteins. PLOS Genetics 11, 1–26 (2015). URL 10.1371/journal.pgen.1005404.

34. Du, W. et al. The 2nd sialic acid-binding site of influenza a virus neuraminidase is an important determinant of the hemagglutinin-neuraminidase-receptor balance. PLoS Pathogens 15, e1007860 (2019).

35. Bedford, T. et al. Global circulation patterns of seasonal influenza viruses vary with antigenic drift. Nature 523, 217–220 (2015).

36. Rubner, Y., Tomasi, C. & Guibas, L. J. A metric for distributions with applications to image databases. In Sixth international conference on computer vision (IEEE Cat. No. 98CH36271), 59–66 (IEEE, 1998).

37. Garten, R. et al. Update: influenza activity in the united states during the 2017–18 season and composition of the 2018–19 influenza vaccine. MMWR. Morbidity and mortality weekly report 67 (2018).

38. Liu, H. et al. Effect of human h3n2 influenza virus reassortment on influenza incidence and severity during the 2017–18 influenza season in the usa: a retrospective observational genomic analysis. The Lancet Microbe 5 (2024).

39. Lee, J. M. et al. Deep mutational scanning of hemagglutinin helps predict evolutionary fates of human h3n2 influenza variants. Proceedings of the National Academy of Sciences 115, E8276–E8285 (2018).

40. Rasmussen, D. A. & Stadler, T. Coupling adaptive molecular evolution to phylodynamics using fitness-dependent birth-death models. eLife 8, e45562 (2019). URL 10.7554/eLife.45562.

41. Bloom, J. D., Gong, L. I. & Baltimore, D. Permissive secondary mutations enable the evolution of influenza oseltamivir resistance. Science 328, 1272–1275 (2010). URL https://www.science.org/doi/abs/10.1126/science.1187816. https://www.science.org/doi/pdf/10.1126/science.1187816.

42. Gong, L. I., Suchard, M. A. & Bloom, J. D. Stability-mediated epistasis constrains the evolution of an influenza protein. eLife 2, e00631 (2013). URL 10.7554/eLife.00631.

43. Strelkowa, N. & Lässig, M. Clonal interference in the evolution of influenza. Genetics 192, 671–682 (2012).

44. Illingworth, C. J. & Mustonen, V. Components of selection in the evolution of the influenza virus: Linkage effects beat inherent selection. PLoS Pathogens 8, e1003091 (2012).

45. Bhatt, S., Holmes, E. C. & Pybus, O. G. The genomic rate of molecular adaptation of the human influenza a virus. Mol Biol Evol 28, 2443–2451 (2011).

46. Meyer, A. G. & Wilke, C. O. Geometric constraints dominate the antigenic evolution of influenza h3n2 hemagglutinin. PLoS Pathogens 11, e1004940 (2015).

47. Thyagarajan, B. & Bloom, J. D. The inherent mutational tolerance and antigenic evolvability of influenza hemagglutinin. elife 3, e03300 (2014).

48. Ahn, J. J., Yu, T. C., Dadonaite, B., Radford, C. E. & Bloom, J. D. Influenza hemagglutinin subtypes have different sequence constraints despite sharing extremely similar structures. Virus Evolution 12, veag018 (2026).

49. Yu, T. C. et al. Pleiotropic mutational effects on function and stability constrain the antigenic evolution of influenza haemagglutinin. Nature ecology & evolution 10, 452–466 (2026).

50. Jorquera, P. A. et al. Insights into the antigenic advancement of influenza a (h3n2) viruses, 2011–2018. Scientific reports 9, 2676 (2019).

51. Lee, N. et al. The use of plant lectins to regulate h1n1 influenza a virus receptor binding activity. PLOS ONE 13, 1–21 (2018). URL 10.1371/journal.pone.0195525.

52. Altman, M. O. et al. Human influenza a virus hemagglutinin glycan evolution follows a temporal pattern to a glycan limit. MBio 10, 10–1128 (2019).

53. Rambaut, A. et al. The genomic and epidemiological dynamics of human influenza a virus. Nature 453, 615–619 (2008).

54. Neverov, A. D., Lezhnina, K. V., Kondrashov, A. S. & Bazykin, G. A. Intrasubtype reassortments cause adaptive amino acid replacements in h3n2 influenza genes. PLoS genetics 10, e1004037 (2014).

