## Supplementary Information for "Sequence-structured fitness landscapes predict influenza evolution"

#### Epidemiological model

We model transmission between individual donors and recipients using a time discrete branching process[46]. Each viral variant $a$ that may cause infection is represented by a protein sequence of length $L$:

$$\mathbf{g}^a = \{g_1^a, g_2^a, ...g_L^a\}$$

where alleles $g_i^a$ take values, at each locus $i$, in an alphabet of size $q = 21$, representing the 20 possible amino acids plus a gap.

The transmissibility effect on the fitness of variant $a$ is represented by the function $w_a$, which modulates its effective reproductive number as $R_a = R(1+w_a)$, where R is the effective reproductive number of the reference variant with fitness of one. The parameter $w_a$, with $|w_a| \ll 1$, plays a role analogous to a selection coefficient in population genetics. Then, the number of new individuals $n_a(t+1)$ infected by the variant $a$ at time $t+1$ given $n_a(t)$ individuals with the same variant at time $t$, follows the negative binomial distribution

$$P_{NB+}(n_a(t+1) \mid n_a(t), k, R_a) = \frac{\Gamma(n_a(t)k + n_a(t+1))}{n_a(t+1)!\,\Gamma(n_a(t)k)} \times \left(\frac{k}{k+R_a}\right)^{n_a(t)k} \left(\frac{R_a}{k+R_a}\right)^{n_a(t+1)} \quad \textbf{(3)}$$

where $k$ stand as dispersion parameter and determines super-spreading. In the large-population, weak-selection regime, this branching process admits a diffusion approximation for allele-frequency trajectories, which we use for inference below.

#### Inferring the fitness effects of mutations from evolutionary histories

Building on the epidemiological model described above, we aim to infer the fitness effects of individual mutations from mutant allele frequency trajectories. To this end, we focus on how allele frequencies evolve across successive rounds of infection according to the branching process. For locus $i$ and allele $\alpha$, the allele frequency at time $t$ is:

$$x_{i,\alpha}(t) = \sum_{a=1}^{M} \delta(g_i^a, \alpha)\, y_a(t), \quad \textbf{(4)}$$

where $y_a(t) = n_a(t)/\sum_b n_b(t)$ is the frequency of the variant $a$ in the population and $\delta(x,y)$ is the Kronecker delta. Here, we assume an additive model for variant fitness, with the fitness of variant $a$ given by the sum of selection coefficients $s_{i,\alpha}$ of its constituent alleles:

$$w_a = \sum_{i=1}^{L} \sum_{\alpha=1}^{q} \delta(g_i^a, \alpha)\, s_{i,\alpha}. \quad \textbf{(5)}$$

For small effects of individual mutations on transmission and large number of infected population, allele frequency evolution can be approximated as a diffusion process and described by a Fokker–Planck equation, in which the expected change in allele frequencies is captured by the drift vector:

$$d_{i,\alpha}(t) = x_{i,\alpha}(t)(1 - x_{i,\alpha}(t))\, s_{i,\alpha} + \sum_{j \neq i, \beta} \big(x_{i\alpha, j\beta}(t) - x_{i,\alpha}(t) x_{j,\beta}(t)\big) s_{j,\beta} \quad \textbf{(6)}$$

and fluctuations in allele frequencies due to the inherent stochasticity are described by the diffusion matrix

$$C_{i\alpha, j\beta}(t) = \left(\frac{1}{k} + \frac{1}{R}\right) \times \begin{cases} x_{i\alpha, j\beta}(t) - x_{i,\alpha}(t) x_{j\beta}(t) & i \neq j \\ x_{i,\alpha}(t)(1 - x_{i,\alpha}(t)) & i = j \end{cases} \quad \textbf{(7)}$$

where the pairwise frequency

$$x_{i\alpha, j\beta}(t) = \sum_{a=1}^{M} \delta\left(g_i^a, \alpha\right) \delta\left(g_j^a, \beta\right) y_a(t) \quad \textbf{(8)}$$

denotes the joint occurrence of alleles $\alpha$ and $\beta$ at loci $i$ and $j$ over time $t$.

The solution of the Fokker–Planck equation via a path-integral approximation yields the two-time transition probability density $P(\mathbf{x}(t_{m+1}) | \mathbf{x}(t_m))$ for the frequency vector $\mathbf{x}(t) = (x_{1,1}(t), x_{1,2}(t), \ldots, x_{1,q}(t), x_{2,1}(t), x_{2,2}(t), \ldots, x_{L,q}(t))$. This allows us to compute the likelihood of an entire evolutionary path of the mutant allele frequencies, $(\mathbf{x}(t_1), \mathbf{x}(t_2), \ldots, \mathbf{x}(t_K))$, given an initial state $\mathbf{x}(t_0)$:

$$P\big(\{\mathbf{x}(t_m)\}_{m=1}^{T} \mid \mathbf{x}(t_0), \mathbf{s}, N\big) \approx \prod_{m=0}^{T-1} \frac{1}{\sqrt{\det \mathbf{C}(\mathbf{x}(t_m))}} \times \left(\frac{N}{2\pi \Delta t_m}\right)^{\frac{qL}{2}} \prod_{i=1}^{qL} dx_i(t_{m+1}) \times \exp\Big(-\frac{N}{2} S\big(\{\mathbf{x}(t)\}_{t=0}^{T}\big)\Big) \quad \textbf{(9)}$$

with $N = \sum_{a=1}^{M} n_a$ is the total number of individuals infected by all variants, $\Delta t_m = t_{m+1} - t_m$ and the action

$$S\big(\{\mathbf{x}(t_m)\}_{m=0}^{T}\big) = \sum_{m=0}^{T-1} \Big[\frac{\mathbf{x}(t_{m+1}) - \mathbf{x}(t_m)}{\Delta t_m} - \mathbf{d}(\mathbf{x}(t_m))\Big]\big(\mathbf{C}^{-1}(\mathbf{x}(t_m))\big[\frac{\mathbf{x}(t_{m+1}) - \mathbf{x}(t_m)}{\Delta t_m} - \mathbf{d}(\mathbf{x}(t_m))\Big] \qquad \textbf{(10)}$$

In practice, our data consists of sets of genetic sequences obtained from a population at multiple time points $t_k$, $k \in \{0, 1, \ldots, K\}$. These sequences can be used to compute the path of mutant allele frequencies as well as the double mutant frequencies defined above. Then, the relationship between observed trajectories and parameters can be inverted by Bayesian inference to obtain the posterior probability of selection coefficients:

$$P\Big(\mathbf{s}|\{\mathbf{x}(t_m)\}_{m=0}^{T}\Big) \propto P\big(\{\mathbf{x}(t_m)\}_{m=1}^{T} \mid \mathbf{x}(t_0), \mathbf{s}, N\big) \times P_{Prior}(\mathbf{s}) \qquad \textbf{(11)}$$

where $P_{Prior}(\mathbf{s})$ is a Gaussian prior probability for the selection coefficients with zero mean and covariance $\frac{1}{\gamma}\mathbf{I}$; precision parameter $\gamma$ serves as a regularization parameter.

The maximum a posteriori estimate of selection coefficients, given an evolutionary history of allele frequencies, is obtained from the Euler–Lagrange equations applied to the posterior. Assuming constant epidemiological parameters {R, N, k} absorbed into $\gamma$, this simplifies to

$$\hat{\mathbf{s}} = [\mathbf{C_{int}} + \gamma \mathbb{I}]^{-1} \mathbf{\Delta x} \qquad \textbf{(12)}$$

where $\mathbf{\Delta x} = \mathbf{x}(t_K) - \mathbf{x}(t_0)$ is the is the net change in allele frequencies, and $\mathbf{C_{int}}$ is the covariance matrix of mutant allele frequencies integrated over time and accounts for competition between variants as well as the speed of growth for different viral lineage.

The posterior distribution of the selection coefficients $\mathbf{s}$ is a multivariate Gaussian distribution, samples of $\mathbf{s}$ are therefore drawn as

$$\mathbf{s} \sim \mathcal{N}\big(\hat{\mathbf{s}}, \mathbf{\Sigma}\big), \qquad \textbf{(13)}$$

where $\hat{\mathbf{s}}$ is the maximum a posteriori estimate from Eq. 12 and $\mathbf{\Sigma}$ is the posterior covariance matrix, given by

$$\mathbf{\Sigma} = \frac{k+R}{NkR} \left(\mathbf{C_{int}} + \gamma \mathbb{I}\right)^{-1} \qquad \textbf{(14)}$$

The diagonal elements of $\mathbf{\Sigma}$ quantify the uncertainty (variance) in the posterior estimate of the selection coefficient $\hat{\mathbf{s}}$.

#### From branching process to Wright–Fisher sampling

For influenza virus, at the seasonal scale, transmission heterogeneity is relatively small, leading to large $k$ values. In the limit $k \to \infty$, the negative binomial distribution for the number of secondary infections converges to a Poisson distribution with mean $R_a$. If each of $N$ infected individuals draws a Poisson distributed number of offspring with mean $R_a$ proportional to its fitness, the total number of offspring across all individuals is Poisson distributed with mean $\Lambda = \sum_{a=1}^{N} R_a$. Conditional on the total number of offspring $K$, the offspring counts per variant follow a multinomial distribution,

$$\mathbf{n} = (n_1, \ldots, n_V) \,\big|\, K \ \sim\ P_{\text{Mult}}(\mathbf{n} \mid K, \mathbf{p}), \quad \text{with } p_a = \frac{R_a}{\Lambda}.$$

This corresponds to a Wright–Fisher sampling step with fitness-proportional reproduction. If the total population size is fixed at $K = N$, the offspring counts are sampled as

$$\mathbf{n} = (n_1, \ldots, n_V) \sim P_{\text{Mult}}(\mathbf{n} \mid N, \mathbf{p}), \quad \mathbf{p} = (p_1, \ldots, p_V).$$

providing a natural Wright–Fisher approximation of the evolutionary dynamics, with selection differences captured by fitness-dependent reproductive numbers.

#### Single-site model

As a reference that ignores genetic linkage, we also consider a single-site model (SSM), in which each locus is treated as an independent population. The SSM differs from the generative model in two respects: selection coefficients are inferred without the linkage-disequilibrium terms of the covariance matrix, and forward dynamics are simulated separately at each locus.

Neglecting linkage disequilibrium sets the off-diagonal entries of $\mathbf{C_{int}}$ to zero, so Eq. 12 decouples into independent estimates for each allele,

$$\hat{s}^{\text{SS}}_{i,\alpha} = \frac{\Delta x_{i,\alpha}}{[\mathbf{C_{int}}]_{i\alpha,i\alpha} + \gamma}, \qquad [\mathbf{C_{int}}]_{i\alpha,i\alpha} = \int x_{i,\alpha}(t)\big(1 - x_{i,\alpha}(t)\big)\, dt, \qquad \textbf{(15)}$$

where the integral runs over the training window. This is the maximum a posteriori estimate for a locus evolving in isolation, under the same Gaussian prior as the full model. In the full model, the off-diagonal terms separate the effect of an allele from that of the alleles linked to it. Without them, $\hat{s}^{\mathrm{SL}}_{i,\alpha}$ attributes to each allele the entire frequency change it experienced, including change driven by selection on linked alleles (genetic hitchhiking).

Forward simulations under the SSM apply the Wright–Fisher step described above to each locus independently, with alleles in place of variants and allele fitness $1+s_{i,\alpha}$ in place of variant fitness $1+w_a$. Starting from the allele frequencies $x_{i,\alpha}(t_0)$ of the population at $t_0$, the allele counts $\mathbf{n}_i = (n_{i,1},\ldots,n_{i,q})$ at locus $i$ are updated each generation as

$$\mathbf{n}_i(t+1) \sim P_{\mathrm{Mult}}\big(\mathbf{n} \mid N, \mathbf{p}_i(t)\big), \qquad p_{i,\alpha}(t) = \frac{x_{i,\alpha}(t)\,(1+s_{i,\alpha})}{\sum_{\beta=1}^{q} x_{i,\beta}(t)\,(1+s_{i,\beta})}, \tag{16}$$

with $x_{i,\alpha}(t+1) = n_{i,\alpha}(t+1)/N$.

The SSM therefore describes a population of unlinked loci. We apply the SSM to the same two tasks as the generative model, described in the following subsections: predicting the fate of individual mutations and forecasting the composition of the population at a fixed horizon.

#### Predicting fixation and loss of mutations

The mutation fate prediction task asks: given a mutation observed at frequency $x$, will it eventually fix or be lost? The quantity of interest is the conditional fixation probability $P(\mathrm{fix} \mid x)$. To estimate it from the data, each mutation observed at a time $t$ with frequency $x(t)$ is treated as one instance, assigned to a frequency bin $B$ according to $x(t)$, and labeled by its later outcome. An instance is labeled fixed (resp. lost) if, following the time point at which its frequency is recorded, the mutation's frequency is above $0.95$ (resp. below $0.05$) for two consecutive observation times; instances that do not meet either criterion are classified as polymorphic and excluded. The empirical fixation probability in bin $B$ is then computed as the fraction of mutations in that bin labeled as fixed:

$$\widehat{P}(\mathrm{fix} \mid x \in B) = n_{fix}(B)/(n_{fix}(B)+n_{lost}(B)). \tag{17}$$

Under neutral drift $P(fix \mid x) = x$, so systematic deviations from this diagonal reflect selection.

To test whether information about a mutation beyond its frequency improves prediction, we estimate the conditional fixation probability $P(\mathrm{fix} \mid x, \phi)$, where $\phi$ is a feature of the mutation evaluated at the same time $t$ as its frequency $x(t)$. Within each frequency bin $B$, instances are split into groups $G$ according to $\phi$ and for each subgroup $(B,G)$, the empirical fixation probability is the fraction of resolved instances labeled fixed:

$$\widehat{P}(\mathrm{fix} \mid x \in B,\ \phi \in G) = \frac{n_{\mathrm{fix}}(B,G)}{n_{\mathrm{fix}}(B,G)+n_{\mathrm{lost}}(B,G)}. \tag{18}$$

Comparing $\widehat{P}$ between groups in the same bin holds the frequency approximately fixed, so a difference between groups reflects information that $\phi$ carries beyond $x$. This conditioning is essential because frequency alone strongly predicts fate. Without it, a feature correlated with $x$ would appear predictive simply because one group tends to sit at higher frequencies, even if $\phi$ did not carry additional information.

##### Generative-model fixation probability (GenFix)

We defined a mutation-level feature using the generative model specified by (i) the likelihood of future evolutionary paths, $P\big(\{\mathbf{x}(t_m)\}_{m=1}^{T} \mid \mathbf{x}(t_0), \mathbf{s}, N\big)$ and (ii) the posterior distribution of selection coefficients inferred via Eq. 11 from all observations up to $t_0$, denoted $\mathcal{D}_{\le t_0}$. Here $t_0$ is the time at which the prediction is made.

Forward predictions are generated as $M$ independent replicates. In replicate $r$, the fitness parameters are sampled as $\mathbf{s}^{(r)} \sim \mathcal{N}(\hat{\mathbf{s}}, \boldsymbol{\Sigma})$, the population is initialized at its observed state at $t_0$, and the variant frequencies are propagated by repeated Wright–Fisher sampling. This samples from the same Markov process that defines the path likelihood: in the Poisson (large-$k$) limit of the negative-binomial branching process, the one-step transition at fixed population size $N$ reduces to Wright–Fisher multinomial sampling. Each replicate runs until absorption, when the population collapses to a single dominant variant.

Let $I_m^{(r)} \in \{0,1\}$ indicate whether the mutation $m$ is carried by the surviving genotype in the replicate $r$. The generative-model fixation probability is

$$\mathrm{GenFix}_m(t_0) = \frac{1}{M}\sum_{r=1}^{M} I_m^{(r)} \tag{19}$$

a Monte Carlo estimate of the probability that mutation $m$ fixes under the model, given the population observed at $t_0$ and averaged over the posterior uncertainty in $\mathbf{s}$:

$$\mathrm{GenFix}_m(t_0) \xrightarrow{M\to\infty} \int P\big(\mathrm{fix}_m \mid \mathbf{x}(t_0), \mathbf{s}, N\big)\, p\big(\mathbf{s} \mid \mathcal{D}_{\le t_0}\big)\, d\mathbf{s}. \tag{20}$$

Because GenFix is computed only from $\mathcal{D}_{\le t_0}$, it uses no information from after the prediction time. To test whether it adds information beyond the observed frequency, we apply the stratification above with $\phi = \mathrm{GenFix}$. For each instance, $\mathrm{GenFix}_m(t_0)$ is computed at the time $t_0$ at which its frequency $x$ is recorded. Instances are pooled across observation times, and within each frequency bin they are split into high- and low-GenFix groups at within-bin quantiles (for example, top vs. bottom half).

##### Single-site fixation probability (SSFix)

As a linkage-free counterpart of GenFix, we compute fixation probabilities under the SSM. For each observation time $t_0$, selection coefficients $\hat{\mathbf{s}}^{\mathrm{SL}}$ are inferred from $\mathcal{D}_{\le t_0}$ (Eq. 15), and each locus is initialized at its allele frequencies at $t_0$. In each of $M$ independent replicates, every locus is propagated with Eq. 16 until absorption, when a single allele remains at that locus. Let $I_{i,\alpha}^{(r)} \in \{0,1\}$ indicate whether allele $\alpha$ is the surviving allele at locus $i$ in replicate $r$. For mutation $m$, corresponding to allele $\alpha$ at locus $i$, the single-site fixation probability is

$$\mathrm{SSFix}_m(t_0) = \frac{1}{M}\sum_{r=1}^{M} I_{i,\alpha}^{(r)}. \tag{21}$$

SSFix differs from GenFix in two ways. Absorption is defined separately at each locus rather than for the whole genotype, so the fate of a mutation depends only on its own allele-frequency trajectory. In addition, selection coefficients are held at $\hat{\mathbf{s}}^{\mathrm{SS}}$ in every replicate rather than sampled from the posterior.

##### Predictive performance with AUROC

Mutations were grouped by their frequency at the observation time $t_0$, into adjacent bins of width $0.1$ and centered at $x \in \{0.1, 0.2, \ldots, 0.9\}$ (implemented as half-open intervals to avoid double counting at bin boundaries). Within each frequency bin, mutations were ranked by $\mathrm{GenFix}_m(t_0)$, by $\mathrm{SLFix}_m(t_0)$, or by their observed frequency (null ranking). Discriminative performance for fixation versus loss was quantified using the receiver operating characteristic (ROC) curve and its area under the curve (AUROC), computed from these scores and the binary outcomes (fix/loss).

To construct the ROC curve, mutations within each bin were sorted from highest to lowest score and we varied the number $k$ of mutations labeled as "predicted to fix" by selecting the top k entries in the ranking (with $k = 0, 1, \ldots, n$ for $n$ mutations in the bin). For each $k$, the resulting predictions define counts of true positives (TP), false positives (FP), true negatives (TN), and false negatives (FN), from which the true positive rate (TPR) and false positive rate (FPR) were computed as

$$\mathrm{TPR} = \frac{\mathrm{TP}}{\mathrm{TP}+\mathrm{FN}}, \qquad \mathrm{FPR} = \frac{\mathrm{FP}}{\mathrm{FP}+\mathrm{TN}}.$$

Each choice of $k$ yields a point (FPR,TPR) on the ROC curve, and AUROC is the area under this curve. AUROC values range from $0$ to $1$, with $1$ indicating perfect classification, $0.5$ random performance, and values below $0.5$ reflecting worse-than-random predictions.

##### Association between fitness statistics and fixation or loss

As described in the main text, we studied how fitness statistics (inferred fitness effect of a mutation, background fitness advantage, and background fitness growth; **Fig. 5A-C**, **Fig. S12-14**) were associated with fixation or loss. We limited our analysis to mutations that were observed at frequencies of >5% for at least 90 days, thereby excluding low-confidence trajectories for mutations that only briefly fluctuated above and below the 5% threshold.

In contrast to predictive tasks, which only used data available up until the prediction time to infer fitness, here we used fitness parameters inferred from the entire trajectory. This is because our goal in these comparisons was not to assess predictive power (which we tested in other ways, as described above), but rather to explore the relationship between (our best estimate of) underlying fitness and evolutionary success. The three statistics that we compared were the fitness effect of the mutation (**Fig. 5A**, **Fig. S12**), the difference in fitness between genotypes carrying the mutation and the population average fitness at the time that the mutation crossed the 5% threshold (**Fig. 5B**, **Fig. S13**), and the overall rate of fitness increase for genotypes carrying the mutation, computed as the difference in the average fitness of genotypes carrying the mutation at the time that it first crossed the 5% threshold and the average fitness of genotypes carrying the mutation at the time that it was subsequently lost (frequency decreased below 5%) or fixed (frequency increased above 95%), divided by the time between when the mutation first rose above 5% and when it was subsequently lost or fixed, measured in years (**Fig. 5C**, **Fig. S14**). These tests involved simple statistical comparisons of the distributions of these quantities between mutations that fixed and those that were lost, and did not involve simulations or forecasts.

#### Forecasting population composition with generative-model simulations

To forecast the future genetic composition of the circulating population, an observation time $t_0$ is fixed and the goal is to predict the population at future time $t_p = t_0 + \Delta$, where $\Delta$ is the forecasting horizon. Model parameters are inferred from training data $\mathcal{D}_{\leq t_0}$ as described above, and forward trajectories are generated by initializing the variant population at the empirical state at $t_0$ and propagating it under Wright–Fisher dynamics with fitness parameters sampled from its posterior distribution. Each simulation yields a predicted population at $t_p$, represented as a discrete distribution over variants with associated frequencies.

To account for stochasticity in the forward dynamics, $M$ independent realizations of the future population are generated, $\{P_1, P_2, \ldots, P_M\}$. A single point forecast $\hat{P}$ is then obtained using a medoid estimator: among the simulated populations, $\hat{P}$ is defined as the population that minimizes the sum of pairwise distances to all other simulated populations,

$$\hat{P} \;=\; \arg \min_{P_i \in \{P_1, \ldots, P_M\}} \sum_{j=1}^{M} \mathrm{EMD}(P_i, P_j), \tag{22}$$

where $\mathrm{EMD}(P_i, P_j)$ is the earth mover's distance between populations $P_i$ and $P_j$ (defined below). Briefly, EMD treats each population as a weighted distribution over sequences and computes the minimum transport cost required to transform one population into the other, using allele-level genetic distance between sequences as the ground metric and variant frequencies as weights. This medoid-based estimator provides a representative forecast that is robust to outlier simulations and stochastic variation across replicates. Forecast accuracy is assessed by comparing $\hat{P}$ to the empirically observed population at $t_p$ using the same EMD metric.

#### Single-site forecasts

Forecasts under the SSM follow the same protocol, with the per-locus dynamics of Eq. 16 run from $t_0$ to $t_p$. Selection coefficients are fixed at $\hat{\mathbf{s}}^{\mathrm{SS}}$, so realizations differ only through genetic drift. Because loci evolve independently, a simulation yields allele frequencies at each locus rather than variant frequencies, and the population at $t_p$ must be reconstructed. We assume linkage equilibrium among loci and restrict the forecast to the variants present at $t_0$, $S_0 = \{a : y_a(t_0) > 0\}$. In realization $r$, each variant $a \in S_0$ receives the weight

$$w_a^{(r)} = \prod_{i=1}^{L} x_{i,g_i^a}^{(r)}(t_p), \qquad \hat{y}_a^{(r)}(t_p) = \frac{w_a^{(r)}}{\sum_{b \in S_0} w_b^{(r)}}, \tag{23}$$

where $x_{i,\alpha}^{(r)}(t_p)$ is the frequency of allele $\alpha$ at locus $i$ at $t_p$ in realization $r$. The product is the probability that a sequence assembled by drawing one allele independently at each locus, according to these frequencies, equals $\mathbf{g}^a$. Allele combinations outside $S_0$ also receive weight under this product. They are discarded because the model includes neither mutation nor recombination that could create them, which keeps the SSM forecast on the same set of variants as the generative model forecast. The distribution $\{\hat{y}_a^{(r)}(t_p)\}_{a \in S_0}$ defines the realization $P_r$, and the point forecast $\hat{P}$ is the medoid of the $M$ realizations, as for the generative model. If no variant in $S_0$ retains nonzero weight in a realization, that is, if every variant has lost at least one of its alleles, that realization is replaced by the population at $t_0$.

#### Earth mover's distance

To quantify differences between populations, we use the earth mover's distance (EMD). Let

$$P_i = \{(s_k^{(i)}, f_k^{(i)})\}_{k=1}^{N_i}, \qquad P_j = \{(s_l^{(j)}, f_l^{(j)})\}_{l=1}^{N_j}$$

denote two populations, where $s_k^{(i)}$ and $s_l^{(j)}$ are sequences observed with frequencies $f_k^{(i)}$ and $f_l^{(j)}$ in populations $P_i$ and $P_j$, respectively. The pairwise dissimilarity between sequences is defined as the Hamming distance

$$d_{kl}^{(i,j)} = \frac{1}{L} \sum_{r=1}^{L} \mathbf{1}\big[s_{k,r}^{(i)} \neq s_{l,r}^{(j)}\big],$$

where $L$ is the sequence length and $\mathbf{1}[\cdot]$ is the indicator function.

We define a non-negative transport matrix $\mathbf{w}^{(i,j)} = \{w_{kl}^{(i,j)}\}$ and the corresponding transport cost

$$C(\mathbf{w}^{(i,j)}) = \sum_{k=1}^{N_i} \sum_{l=1}^{N_j} d_{kl}^{(i,j)} \, w_{kl}^{(i,j)}.$$

The EMD between $P_i$ and $P_j$ is then obtained by solving the optimal transport problem

$$\mathrm{EMD}(P_i, P_j) = \min_{\mathbf{w}^{(i,j)}} C(\mathbf{w}^{(i,j)})$$

subject to

$$\sum_{k=1}^{N_i} w_{kl}^{(i,j)} = f_l^{(j)}, \qquad \sum_{l=1}^{N_j} w_{kl}^{(i,j)} = f_k^{(i)}, \qquad w_{kl}^{(i,j)} \geq 0.$$

This metric represents the minimum total cost required to transform the frequency distribution of $P_i$ into that of $P_j$, where cost reflects the amount of population frequency that must be reassigned between genetically distinct sequences. In this way, the EMD quantifies the minimal average genetic change needed to transform one population into another, providing a natural measure of dissimilarity between variant populations

##### Null model and lower-bound population

To evaluate forecasting performance, we generated two reference populations. The null model assumes that the population at the observation time remains unchanged into the future, with strain frequencies identical to those observed at the current timepoint $t_0$.

The lower-bound population estimates the minimal achievable earth mover's distance given the sequences available at $t_0$. Construction begins by initializing the frequency of each current strain to zero. For each strain observed at the prediction timepoint $t_p$, we then identified its closest match in the current population based on Hamming distance and added the frequency of the future strain to this nearest neighbor. This procedure allows each strain in the current population to accumulate contributions from multiple future strains, producing a population that represents the best possible approximation of the future under the constraint of using only previously observed sequences.

#### Synthetic data

To generate synthetic data, we implemented a superspreading branching process that models the number of new infections across generations. We considered sequences of length $L = 20$ over an alphabet of $q = 4$ residue states. Selection coefficients were independently sampled from a Gaussian distribution with mean zero and standard deviation $\sigma_{\vec{s}} = 0.05$.

The initial population consisted of $N = 1{,}000$ individuals distributed among 10 randomly generated variants. For each infected individual, the number of secondary infections was drawn from a negative binomial distribution with dispersion parameter $k = 0.1$. The effective reproductive number of the reference variant was initially set to $R = 2.0$. Once the population exceeded $10{,}000$ individuals, $R$ was adaptively adjusted to maintain a mean reproductive number close to 1, thereby stabilizing the population around this threshold. Mutations were introduced at each generation at a rate of $\mu = 10^{-3}$, and the population was evolved for $1{,}000$ generations.

#### Influenza A virus data

We analyzed genome sequences of the surface proteins HA and NA from influenza A/H3N2 and A/H1N1 variants available in GISAID, along with metadata providing collection dates corresponding to the time when individuals were sequenced. The analyzed period spans January 1, 2009, to March 1, 2020, for A/H3N2, and May 1, 2009, to March 1, 2020, for A/H1N1. The starting dates were determined by data availability, as the number of monthly sequences was too limited prior to these dates. The endpoint coincides with the onset of the COVID-19 pandemic, which disrupted influenza evolution.

To ensure data quality, we applied the following preprocessing steps. Sequences without a collection date were discarded, as were sequences in which 5% or more of the positions were gaps or ambiguous amino acids. Sequences whose length differed from that of the full-length protein (566 amino acids for HA, 469 for NA) were also removed, and identical sequences collected on the same date were merged into a single entry with a count. Remaining gaps and ambiguous amino acids were then replaced by the most frequent amino acid at the same position. Finally, to remove likely sequencing or annotation errors, we divided the study period into consecutive windows of approximately one year and, within each window, removed sequences that differed at more than 20 positions from the most common sequence on the window's first collection date. After preprocessing, the dataset comprised $57{,}638$ HA and $53{,}137$ NA sequences for A/H3N2, and $52{,}660$ HA and $45{,}402$ NA sequences for A/H1N1.

##### Smoothing trajectories

As sequences are unevenly distributed over time, accurately estimating frequencies and covariances is challenging: standard binning approaches often produce bins with very few sequences, leading to biased estimates. To address this, we apply a smoothing procedure that uses the exact collection dates of sequences and weights them with a Gaussian factor.

The smoothing process involves stepping through time in discrete intervals of length $t_s$. At each time point $t$, only sequences with collection times $t'$ satisfying $|t - t'| < t_{\text{cutoff}}$ were included. Each sequence count is then weighted using a Gaussian factor proportional to $\exp\left(-\frac{(t-t')^2}{2\tau^2}\right)$, where $\tau$ is a parameter that controls the degree of smoothing. This Gaussian weighting helps to emphasize sequences collected closer in time to $t$ while down-weighting more distant observations. Weighted sequences were then used to compute frequencies and covariances at each time step, producing a continuous representation of the trajectories and reducing artificial fluctuations arising from discrete binning. A uniform smoothing scheme with parameters $t_s = 10$ days, $\tau = 60$ days and $t_{\text{cutoff}} = 3\tau$ was applied consistently in all analyzes.

To prevent data leakage in predictive analyzes, sequences corresponding to testing intervals were excluded from the smoothing procedure. Consequently, the frequency of observing an amino acid $\alpha$ in position $i$ at time $t_0$: $x_{i,\alpha}(t_0)$ results from the Gaussian weighted average of sequences collected within the interval $[t_i : t_0]$.

#### Calculating frequency changes and covariances

To increase robustness to finite sampling in time, we integrated terms in Eq. (10) over time by assuming that frequencies are piecewise linear, rather than summing contributions from each time point. This results in diagonal terms of the integrated covariance given by

$$\sum_{m=0}^{T-1} \Delta t_m \left[ \frac{(3-2x_i(t_{m+1}))(x_i(t_m)+x_i(t_{m+1}))}{6} - \frac{x_i^2(t_m)}{3} \right] \tag{24}$$

and off-diagonal elements

$$\sum_{m=0}^{T-1} \Delta t_m \left[ \frac{x_{ij}(t_m)+x_{ij}(t_{m+1})}{2} - \frac{x_i(t_m)x_j(t_m)+x_i(t_{m+1})x_j(t_{m+1})}{3} - \frac{x_i(t_m)x_j(t_{m+1})+x_i(t_{m+1})x_j(t_m)}{6} \right] \tag{25}$$

#### Choice of regularization

The regularization strength $\gamma$ controls the width of the prior distribution for amino acid residue selection coefficients and therefore affects both the inferred coefficients and their associated uncertainty. Regularization reduces the noise in the inferred coefficients that arise from finite sampling of viral sequences. This is particularly important for amino acid variants observed only transiently in the time-series data, whose integrated covariance matrix in the denominator of Eq. 12 is small.

We selected $\gamma$ using two strategies, depending on the task. For inference of subtype fitness landscapes and fixation forecasting, we used an unsupervised stability criterion. We inferred selection coefficients across a range of regularization strengths ($10 \leq \gamma \leq 60$) and examined pairwise correlations between the resulting estimates (see Fig. 19). The estimates became stable at $\gamma \geq 20$, with strong pairwise correlations ($R^2 > 0.95$) and highly consistent rank orderings. We therefore selected $\gamma = 20$, the smallest value within this stability plateau, to suppress noise while limiting the shrinkage of the inferred coefficients.

For forecasting future viral populations, we jointly selected $\gamma$ and a variant frequency threshold, below which predicted variants were removed from the forecast population, using expanding-window walk-forward cross-validation. For each held-out test date and forecast horizon of $\Delta$ days, the training set included only earlier forecast dates for which the target population, observed $\Delta$ days later, was available on or before the test date. This restriction prevented leakage of future information into hyperparameter selection. We evaluated candidate combinations of $\gamma$ and the frequency threshold on these historical forecasts and selected the combination with the lowest mean training Earth mover's distance (EMD). We then used the selected parameters to generate the forecast for the held-out test date (see Fig. 6).

#### Calculating selection coefficients

After preprocessing, we computed selection coefficients for each subtype and surface protein using Eq. 12. To exclude extremely rare variants and potential sequencing errors, we retained only the selection coefficients of mutations whose frequency exceeded $1\%$ at two or more time points within the training window. For each training window, selection coefficients were expressed relative to a reference allele at each locus, taken as the most frequent allele at the first time point of the window (that is, the consensus sequence at that time), so that $\hat{s}_{i,\mathrm{ref}(i)} = 0$.

### Multinomial logistic regression

Operational forecasts of viral variant frequencies are dominated by multinomial logistic regression (MLR) and related models, which treat the log-odds of each variant as approximately linear in time. These models yield interpretable growth advantages and short-horizon frequency forecasts; public tools such as Nextstrain use MLR on daily sequence counts to estimate variant frequencies and growth advantages[47].

We implemented MLR as a baseline for forecasting variant composition. At each prediction time $t_0$, the sequences observed up to $t_0$ were collapsed into $V$ unique variants, and each variant was assigned smoothed effective counts $Y_{t,v}$ at the grid times $t$, using the Gaussian smoothing of our main method. The MLR models variant probabilities as a softmax of logits that are linear in time,

$$p_v(t) = \frac{\exp(\alpha_v + \beta_v \tilde{t})}{\sum_{u=1}^{V} \exp(\alpha_u + \beta_u \tilde{t})}, \qquad \tilde{t} = \frac{t - \bar{t}}{s_t}, \tag{26}$$

where $\bar{t}$ and $s_t$ are the mean and standard deviation of the training grid times; standardizing time improves numerical conditioning. Because the softmax is unchanged when the same constant is added to every logit, the parameters are identified by fixing the intercept and slope of one variant $r$ to zero, $\alpha_r = \beta_r = 0$. Any variant can serve as a reference; we used the first variant in the list of unique sequences.

The parameters were estimated by minimizing the multinomial negative log-likelihood of the smoothed counts with a ridge penalty on the deviations of the slopes from their mean,

$$J(\boldsymbol{\alpha},\boldsymbol{\beta}) = -\sum_t \sum_{v=1}^{V} Y_{t,v} \log p_v(t) + \lambda \sum_{v=1}^{V} \left(\beta_v - \bar{\beta}\right)^2, \qquad \bar{\beta} = \frac{1}{V}\sum_{v=1}^{V} \beta_v, \tag{27}$$

where $\bar{\beta}$ is computed from the slopes being fitted rather than fixed in advance. We used $\lambda = 0.1$ and minimized $J$ with L-BFGS and a backtracking line search (at most 20,000 iterations, gradient tolerance $10^{-6}$). Because both terms are unchanged when the same constant is added to every slope, the fitted forecasts do not depend on the choice of $r$, which serves only to make the parameters identifiable.

For forecasting, probabilities at a future time $t_p$ were computed with the same standardization, $\tilde{t}_p = (t_p - \bar{t})/s_t$. The predictions were restricted to variants with a nonzero smoothed count at $t_0$, and their probabilities were renormalized to sum to one.

#### Cross-subtype comparison of selection profiles via reference-based mapping

To compare site-specific selection coefficients between H1 (N1) and H3 (N2), we defined a common coordinate system based on an H3 (N2) reference sequence and used it to express both subtype-specific selection profiles on the same positional axis. First, we generated a cross-subtype H1–H3 (N1–N2) alignment using MAFFT profile–profile alignment, starting from subtype-specific alignments for H1 (N1) and H3 (N2). From the merged alignment, we extracted one representative H1 (N1) sequence row and the H3 (N2) reference row and converted their aligned representation into a column-wise mapping table using a gap-aware parser. For each alignment column, this procedure assigned an H1 (N1) residue index (or missing if the H1 sequence contained a gap) and an H3 (N2) reference residue index (or missing if the H3 reference contained a gap). Columns with non-missing indices on both sides define a direct correspondence between H1 (N1) and H3 (N2) reference positions.

We then used this correspondence to express the H1 (N1) selection profile in H3 (N2) coordinates. Specifically, selection coefficients estimated on the H1 (N1) alignment were re-expressed in H3 (N2) coordinates by assigning each H3 (N2) position the selection coefficient of its mapped homologous H1 (N1) position (excluding columns where the mapping was undefined). This yielded two selection vectors indexed by H3 (N2) reference position: an H3 (N2) vector $\hat{\mathbf{s}}_{\mathrm{H3}}$ ($\hat{\mathbf{s}}_{\mathrm{N2}}$) and an H1 (N1)-derived vector $\hat{\mathbf{s}}_{\mathrm{H1}}$ ($\hat{\mathbf{s}}_{\mathrm{N1}}$) in H3 (N2) coordinates. In subsequent analysis, missing values were handled explicitly, and the vectors were smoothed using a Gaussian kernel to reduce noise while preserving broad spatial patterns along the protein. Comparison of the full site-specific selection profiles across subtypes showed a stronger overall correlation for $\hat{\mathbf{s}}_{\mathrm{H1}}$–$\hat{\mathbf{s}}_{\mathrm{H3}}$, consistent with more similar selection landscapes, whereas $\hat{\mathbf{s}}_{\mathrm{N1}}$–$\hat{\mathbf{s}}_{\mathrm{N2}}$ exhibited a weaker overall correlation, suggesting more distinct selection landscapes.

#### Data and code

Raw data and code used in our analysis is available in the GitHub repository located at https://github.com/bartonlab/paper-influenza-prediction. This repository also contains Jupyter notebooks that can be run to reproduce the results presented here.

### References


46. Lee, B. *et al.* Inferring effects of mutations on sars-cov-2 transmission from genomic surveillance data. *Nature Communications* **16**, 441 (2025).
47. Abousamra, E., Figgins, M. & Bedford, T. Fitness models provide accurate short-term forecasts of sars-cov-2 variant frequency. *PLOS Computational Biology* **20**, e1012443 (2024).
48. Jorquera, P. A. *et al.* Insights into the antigenic advancement of influenza a (h3n2) viruses, 2011–2018. *Scientific reports* **9**, 2676 (2019).
49. Lee, J. M. *et al.* Mapping person-to-person variation in viral mutations that escape polyclonal serum targeting influenza hemagglutinin. *eLife* **8**, e49324 (2019).
50. Wiley, D. C., Wilson, I. A. & Skehel, J. J. Structural identification of the antibody-binding sites of hong kong influenza haemagglutinin and their involvement in antigenic variation. *Nature* **289**, 373–378 (1981). URL https://doi.org/10.1038/289373a0.
51. Lee MS, C. J. Predicting antigenic variants of influenza a/h3n2 viruses. *Emerging infectious diseases* **8**, 1385–90 (2004).
52. Koel, B. F. *et al.* Substitutions near the receptor binding site determine major antigenic change during influenza virus evolution. *Science* **342**, 976–979 (2013).
53. Beer, K. *et al.* Characterization of neutralizing epitopes in antigenic site b of recently circulating influenza a(h3n2) viruses. *Journal of General Virology* **99**, 1001–1011 (2018).
54. Chambers, B. S., Parkhouse, K., Ross, T. M., Alby, K. & Hensley, S. E. Identification of hemagglutinin residues responsible for h3n2 antigenic drift during the 2014–2015 influenza season. *Cell Reports* **12**, 1–6 (2015).
55. Zost, S. J. *et al.* Identification of antibodies targeting the h3n2 hemagglutinin receptor binding site following vaccination of humans. *Cell Reports* **29**, 4460–4470.e8 (2019).
56. Popova, L. *et al.* Immunodominance of antigenic site b over site a of hemagglutinin of recent h3n2 influenza viruses. *PLoS ONE* **7**, e41895 (2012).
57. Iba, Y. *et al.* Conserved neutralizing epitope at globular head of hemagglutinin in h3n2 influenza viruses. *Journal of Virology* **88**, 7130–7144 (2014). URL https://journals.asm.org/doi/abs/10.1128/jvi.00420-14. https://journals.asm.org/doi/pdf/10.1128/jvi.00420-14.
58. Weis, W. *et al.* Structure of the influenza virus haemagglutinin complexed with its receptor, sialic acid. *Nature* **333**, 426–431 (1988). URL https://doi.org/10.1038/333426a0.
59. Yang, H. *et al.* Structure and receptor binding preferences of recombinant human a(h3n2) virus hemagglutinins. *Virology* **477**, 18–31 (2015). URL https://www.sciencedirect.com/science/article/pii/S004268221400573X.
60. Altman, M. O. *et al.* Human influenza a virus hemagglutinin glycan evolution follows a temporal pattern to a glycan limit. *MBio* **10**, 10–1128 (2019).
61. Caton, A. J., Brownlee, G. G., Yewdell, J. W. & Gerhard, W. The antigenic structure of the influenza virus a/pr/8/34 hemagglutinin (h1 subtype). *Cell* **31**, 417–427 (1982).
62. Matsuzaki, Y. *et al.* Epitope mapping of the hemagglutinin molecule of a/(h1n1)pdm09 influenza virus by using monoclonal antibody escape mutants. *Journal of Virology* **88**, 12364–12373 (2014).

63. Xu, R. *et al.* Structural basis of preexisting immunity to the 2009 h1n1 pandemic influenza virus. *Science* **328**, 357–360 (2010).
64. Linderman, S. L. *et al.* Potential antigenic explanation for atypical h1n1 infections among middle-aged adults during the 2013–2014 influenza season. *Proceedings of the National Academy of Sciences* **111**, 15798–15803 (2014).
65. Guthmiller, J. J. *et al.* First exposure to the pandemic h1n1 virus induced broadly neutralizing antibodies targeting hemagglutinin head epitopes. *Science Translational Medicine* **13**, eabg4535 (2021).
66. Yasuhara, A. *et al.* Antigenic drift originating from changes to the lateral surface of the neuraminidase head of influenza a virus. *Nature Microbiology* **4**, 1024–1034 (2019). URL https://doi.org/10.1038/s41564-019-0401-1.
67. Lee, N. *et al.* The use of plant lectins to regulate h1n1 influenza a virus receptor binding activity. *PLOS ONE* **13**, 1–21 (2018). URL https://doi.org/10.1371/journal.pone.0195525.
68. Thompson, C. P. *et al.* A naturally protective epitope of limited variability as an influenza vaccine target. *Nature Communications* **9**, 3859 (2018).
69. Immune Epitope Database. H1n1 ha peptide assay 22971621: Rsqlknnak. https://www.iedb.org/assay/22971621. Accessed 2026.
70. Yasugi, M. *et al.* Emerging antigenic variants at the antigenic site sb in pandemic a(h1n1)2009 influenza virus in japan detected by a human monoclonal antibody. *PLoS ONE* **8**, e77892 (2013).
71. Lee, J. M. *et al.* Deep mutational scanning of hemagglutinin helps predict evolutionary fates of human h3n2 influenza variants. *Proceedings of the National Academy of Sciences* **115**, E8276–E8285 (2018).
72. Ketklao, S., Boonarkart, C., Phakaratsakul, S., Auewarakul, P. & Suptawiwat, O. Responses to the sb epitope contributed to antigenic drift of the influenza a 2009 h1n1 virus. *Archives of Virology* **165**, 2503–2512 (2020).
73. Ge, J. *et al.* The antibody response against neuraminidase in human influenza a (h3n2) virus infections during 2018/2019 flu season: Focusing on the epitopes of 329-n-glycosylation and e344 in n2. *Frontiers in Microbiology* **13**, 845088 (2022).
74. Wang, Y., Lei, R., Nourmohammad, A. & Wu, N. C. Antigenic evolution of human influenza h3n2 neuraminidase is constrained by charge balancing. *eLife* **10**, e72516 (2021). URL https://doi.org/10.7554/eLife.72516.
75. Jo, G. *et al.* Structural basis of broad protection against influenza virus by human antibodies targeting the neuraminidase active site via a recurring motif in cdr h3. *Nature Communications* **16**, 7067 (2025). URL https://doi.org/10.1038/s41467-025-62174-2.
76. Air, G., Els, M., Brown, L., Laver, W. & Webster, R. Location of antigenic sites on the three-dimensional structure of the influenza n2 virus neuraminidase. *Virology* **145**, 237–248 (1985). URL https://www.sciencedirect.com/science/article/pii/0042682285901576.
77. Colman, P. M., Varghese, J. N. & Laver, W. G. Structure of the catalytic and antigenic sites in influenza virus neuraminidase. *Nature* **303**, 41–44 (1983). URL https://doi.org/10.1038/303041a0.
78. Wan, H. *et al.* The neuraminidase of a(h3n2) influenza viruses circulating since 2016 is antigenically distinct from the a/hong kong/4801/2014 vaccine strain. *Nature Microbiology* **4**, 2216–2225 (2019).
79. Du, W. *et al.* The 2nd sialic acid-binding site of influenza a virus neuraminidase is an important determinant of the hemagglutinin-neuraminidase-receptor balance. *PLoS Pathogens* **15**, e1007860 (2019).
80. Doud, M. B., Hensley, S. E. & Bloom, J. D. Complete mapping of viral escape from neutralizing antibodies. *PLoS Pathogens* **13**, e1006271 (2017).
81. Gulati, U. *et al.* Antibody epitopes on the neuraminidase of a recent h3n2 influenza virus (a/memphis/31/98). *Journal of Virology* **76**, 12274–12280 (2002). URL https://journals.asm.org/doi/abs/10.1128/jvi.76.23.12274-12280.2002. https://journals.asm.org/doi/pdf/10.1128/jvi.76.23.12274-12280.2002.
82. Lee, J. T. & Air, G. M. Interaction between a 1998 human influenza virus n2 neuraminidase and monoclonal antibody mem5. *Virology* **345**, 424–433 (2006).
83. Dai, M. *et al.* Analysis of the evolution of pandemic influenza a (h1n1) virus neuraminidase reveals entanglement of different phenotypic characteristics. *MBio* **12**, 10–1128 (2021).
84. Wan, H. *et al.* Molecular basis for broad neuraminidase immunity: Conserved epitopes in seasonal and pandemic h1n1 as well as h5n1 influenza viruses. *Journal of Virology* **87**, 9290–9300 (2013).
85. Wan, H. *et al.* Structural characterization of a protective epitope spanning a(h1n1)pdm09 influenza virus neuraminidase monomers. *Nature Communications* **6**, 6114 (2015).
86. Gao, J. *et al.* Antigenic drift of the influenza a(h1n1)pdm09 virus neuraminidase results in reduced effectiveness of a/california/7/2009 (h1n1pdm09)-specific antibodies. *mBio* **10**, e00307–19 (2019).
87. Du, W. *et al.* Substrate binding by the second sialic acid-binding site of influenza a virus n1 neuraminidase contributes to enzymatic activity. *Journal of Virology* **92**, 10.1128/jvi.01243–18 (2018). URL https://journals.asm.org/doi/abs/10.1128/jvi.01243-18. https://journals.asm.org/doi/pdf/10.1128/jvi.01243-18.
88. Catani, J. P. P. *et al.* The antigenic landscape of n1 neuraminidase in human influenza a virus strains isolated between 2009 and 2020. *eLife Reviewed Preprint* (2025).
89. Kirkpatrick Roubidoux, E. *et al.* Novel epitopes of the influenza virus n1 neuraminidase targeted by human monoclonal antibodies. *Journal of Virology* **96**, e00332–22 (2022).

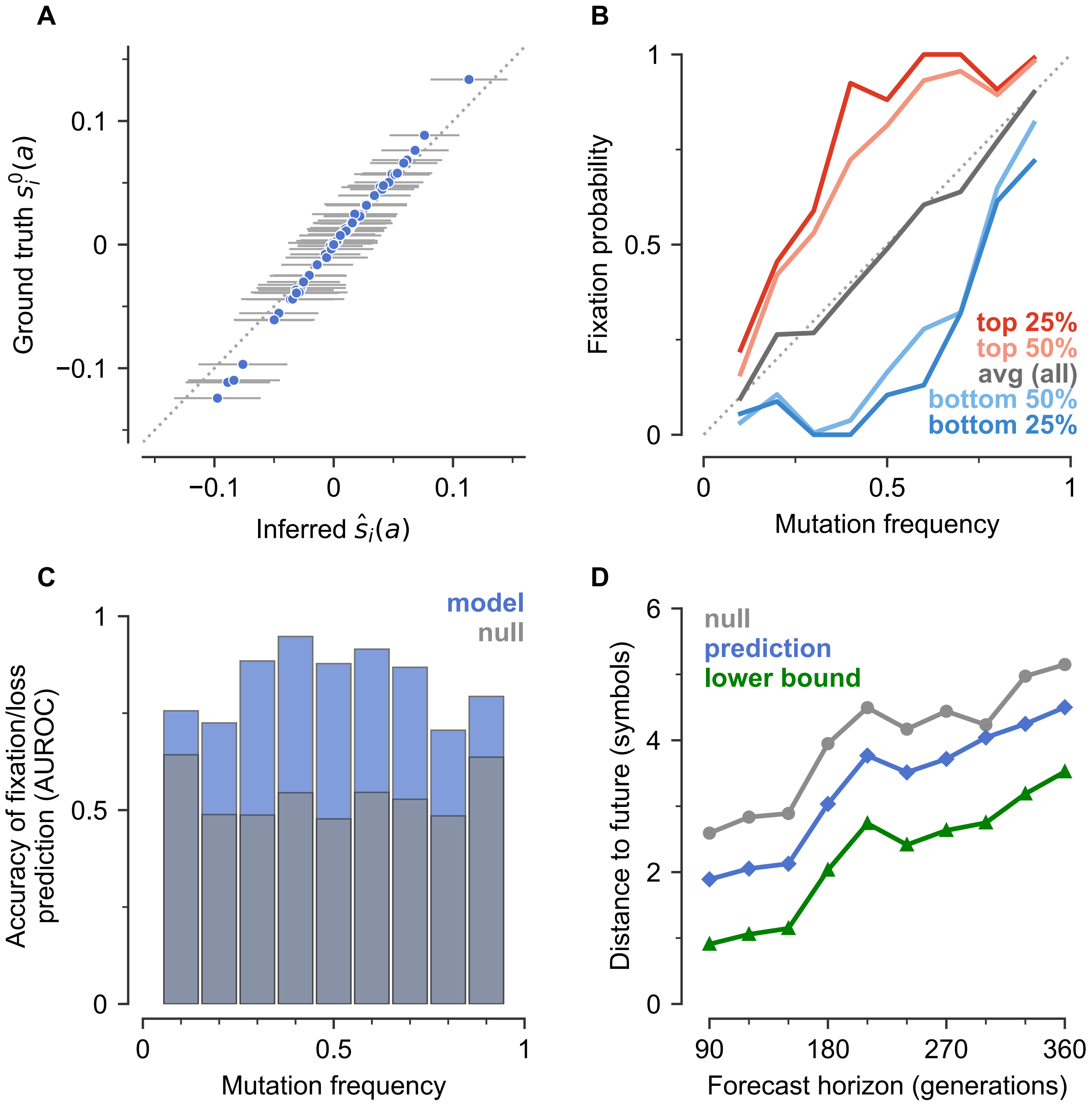


**Fig. S1. Inference method accurately estimates transmission effects of mutations and forecasts evolution in simulations.** Simulated epidemiological dynamics beginning with a mixed population containing variants with beneficial, neutral, and deleterious mutations ($L = 20$ sites, $q = 4$ states per site). **A**, Inferred selection coefficients. Tests on 1000 replicate simulations with identical parameters show that inferred selection coefficients are centered around their true values. Points show the mean inferred value for each mutation and horizontal bars one standard deviation across replicates. **B**, Probability that a mutation fixes as a function of its frequency when the prediction is made. Mutations are ranked by the model's predicted fixation score: top 25% and top 50% (dark and light red), bottom 50% and bottom 25% (light and dark blue), and all mutations (gray). The dotted line marks the neutral expectation, in which a mutation's fixation probability equals its frequency. Mutations ranked highly by the model fix more often than this expectation, and low-ranked mutations are more often lost. **C**, Accuracy of distinguishing mutations that fix from those that are lost (AUROC) as a function of mutation frequency. The model (blue) is compared with a null model that ranks mutations by their current frequency (gray, overlaid). The model outperforms the null model at every frequency. **D**, Earth mover's distance (EMD) between predicted and true population distributions as a function of forecast horizon (generations after the observation window of 300 generations). Predictions from the learned model (blue) are compared to a null model (gray), and to a lower bound representing the best possible estimate given the observed data (green). Distances are measured in symbol units, where each sequence position can take one of $q = 4$ possible states.

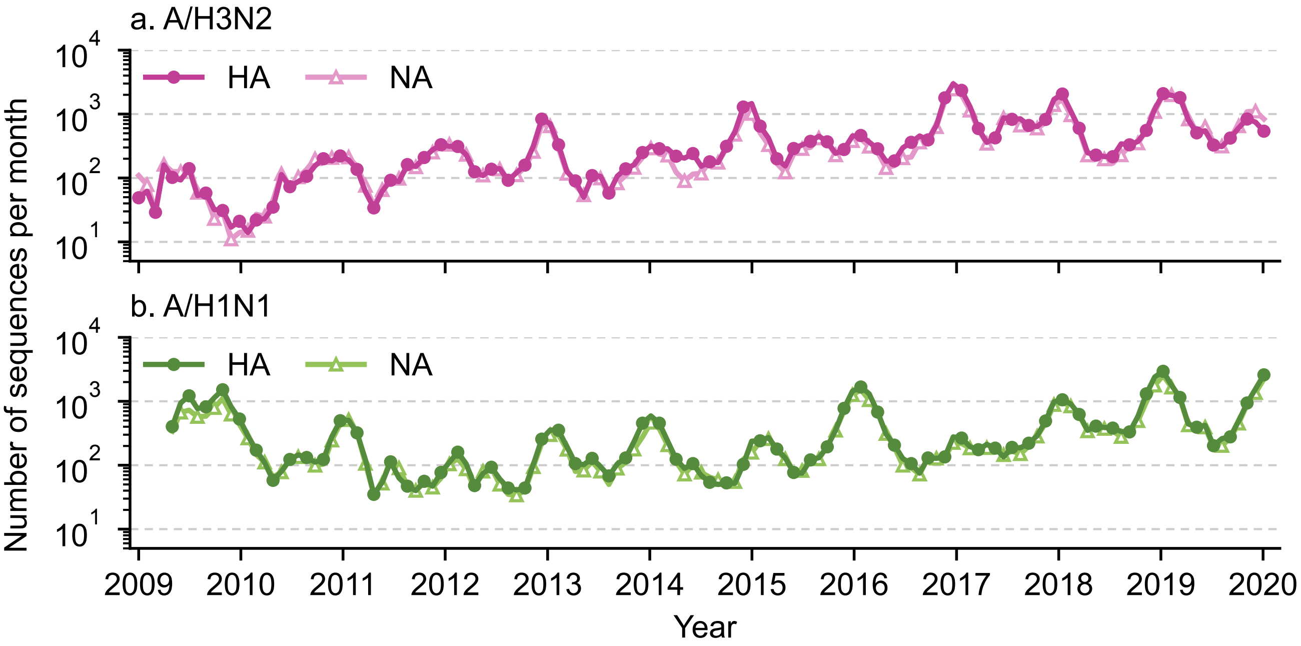


**Fig. S2. Number of HA and NA protein sequences for the influenza A/H3N2 and A/H1N1 subtypes obtained from GISAID.** The analysis covered the period from January 1, 2009, to March 1, 2020, for A/H3N2 and from May 1, 2009, to March 1, 2020, for A/H1N1.

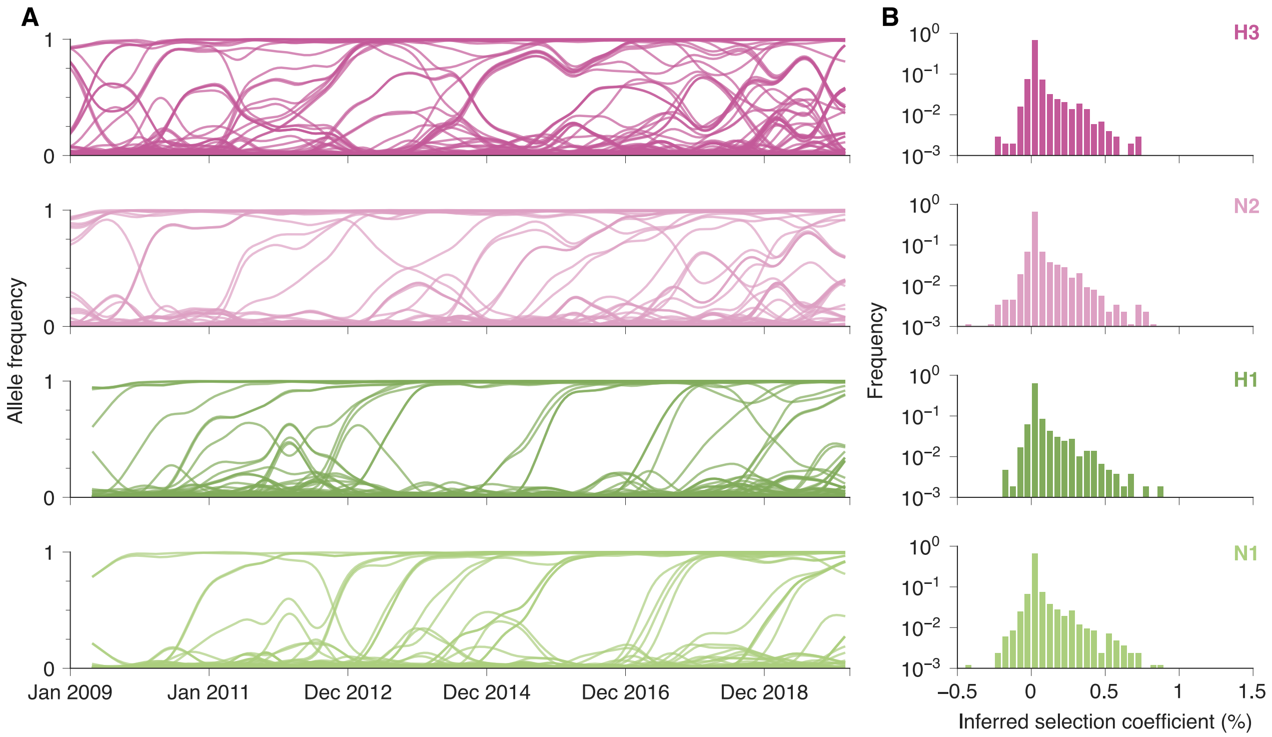


**Fig. S3. Patterns of IAV evolution and selection.** (**A**) Trajectories of mutant allele frequencies over the observation period in H3, N2. H1, and N1. (**B**) Inferred selection coefficients for IAV mutations. A similar pattern appears for each surface protein and subtype, featuring a long tail of beneficial mutations, a large peak near neutrality, and a sharp tail of deleterious mutations.

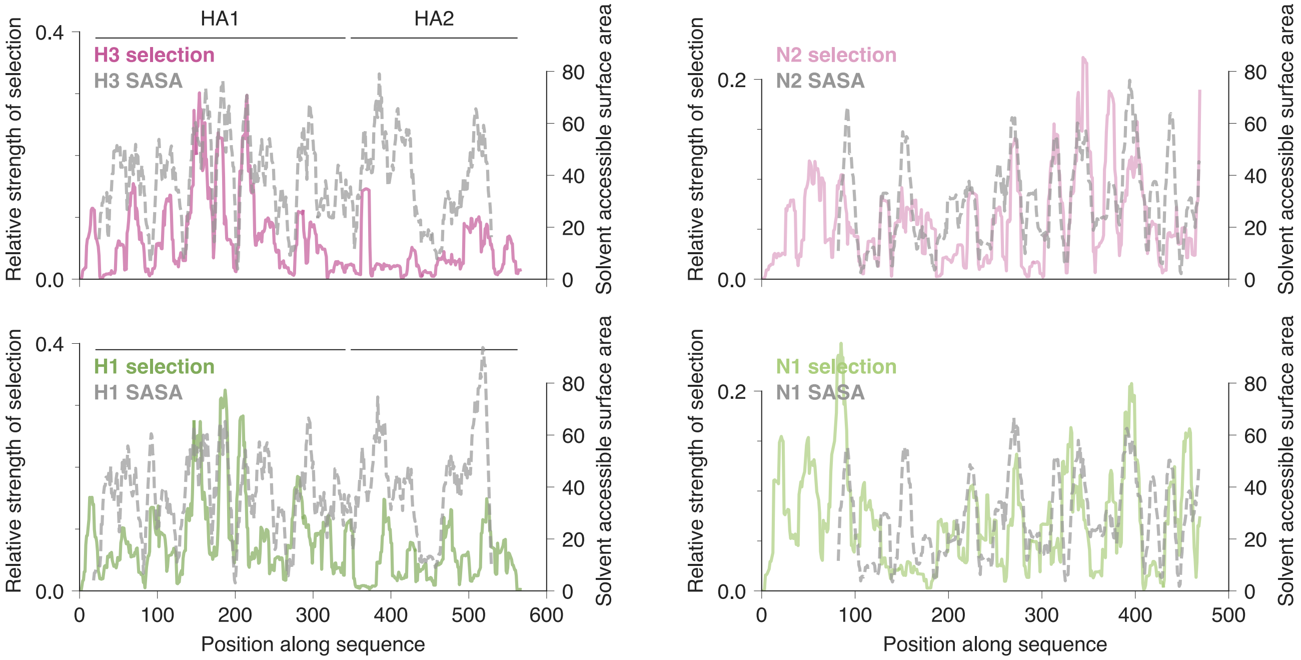


**Fig. S4. Selection versus solvent accessible surface area (SASA).** Patterns of inferred selection and SASA, both smoothed along the protein sequence (Methods), are correlated. We observe the strongest correlations in HA1, followed by NA, with the weakest in HA2. Overall, the correlation between inferred selection and SASA is stronger for H3N2 than for H1N1.

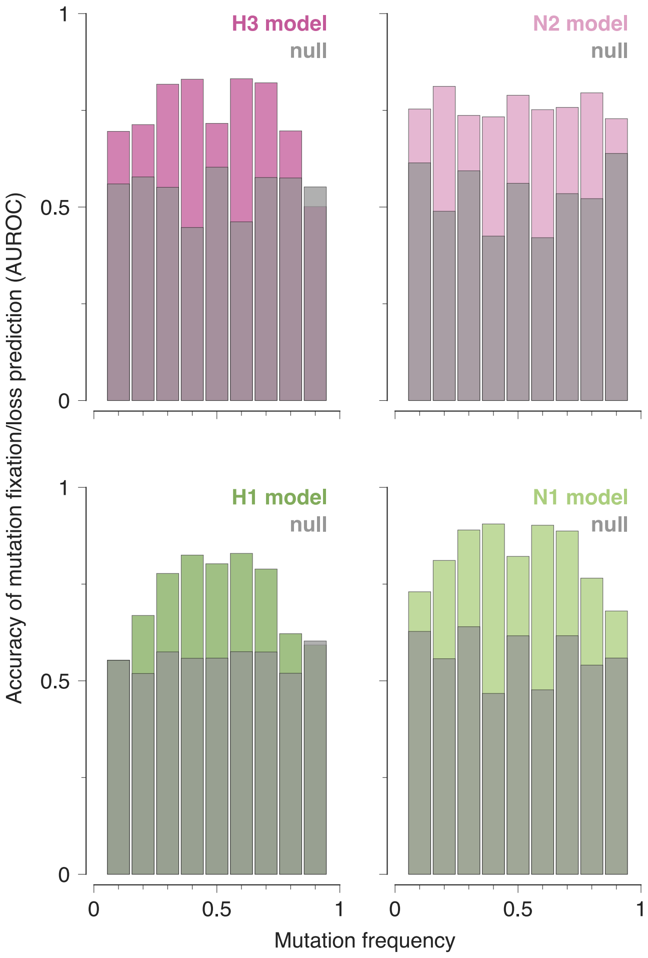


**Fig. S5. Accuracy of predictions for IAV mutation fixation or loss.** Predictions from the sequence-structured model and a neutral model are compared using the area under the receiver operating characteristic curve (AUROC) in each mutation frequency bin. The sequence-structured model consistently exceeds neutral predictions, with the largest improvements observed for mutations at intermediate frequencies where evolutionary outcomes remain most uncertain.

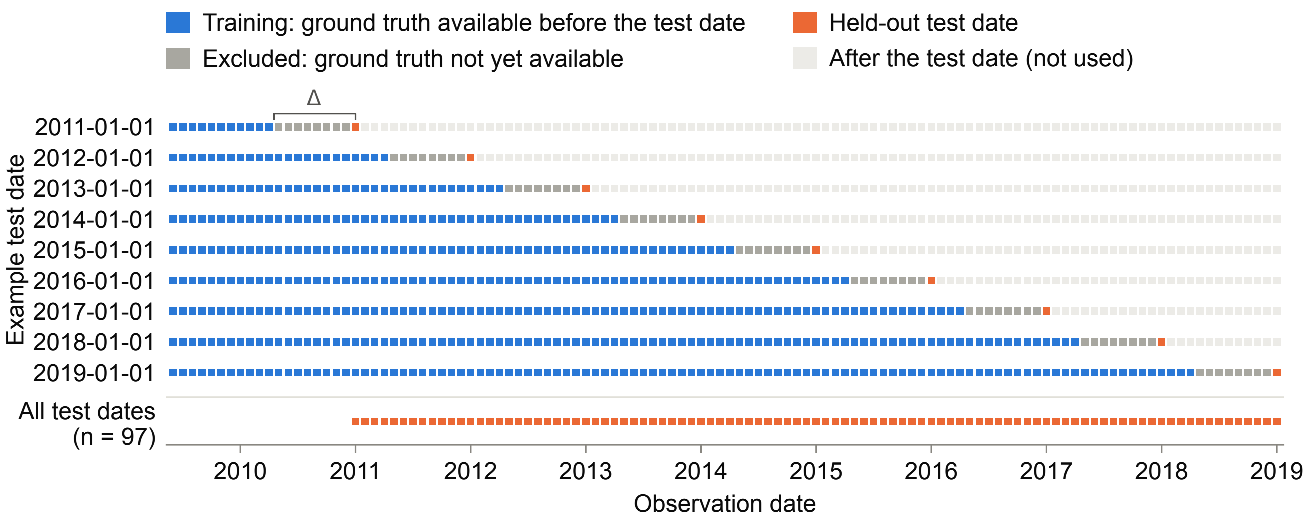


**Fig. S6. Rolling-origin cross-validation with a prediction horizon of 9 months.** Each square is one monthly observation date. The upper rows show the training–test split for example test dates (1 January of each year, 2011–2019); they are illustrative only. The evaluation itself is repeated for every test date marked in the bottom row: the first day of every month from 2011-01-01 through 2019-01-01, inclusive (97 test dates). For each held-out test date $t_0$, the training split (blue) contains only earlier observations whose ground truth, resolved $\Delta$ days after the observation date, was available on or before $t_0$ (observation date $+ \Delta \leq t_0$). Earlier observations whose ground truth resolves after $t_0$ (gray) are excluded, and observations after $t_0$ (light gray) are not used; this restriction prevents information leakage. Within the eligible training split, the combination of $\gamma$ and frequency threshold with the lowest mean training Earth Mover's Distance (EMD) is selected from 21 candidate combinations ($\gamma \in \{20, 40, 100, 150, 200, 250, 300\}$; frequency threshold $\in \{10^{-4}, 10^{-3}, 10^{-2}\}$), and the selected parameters are then used to generate the prediction for the held-out test date (orange).

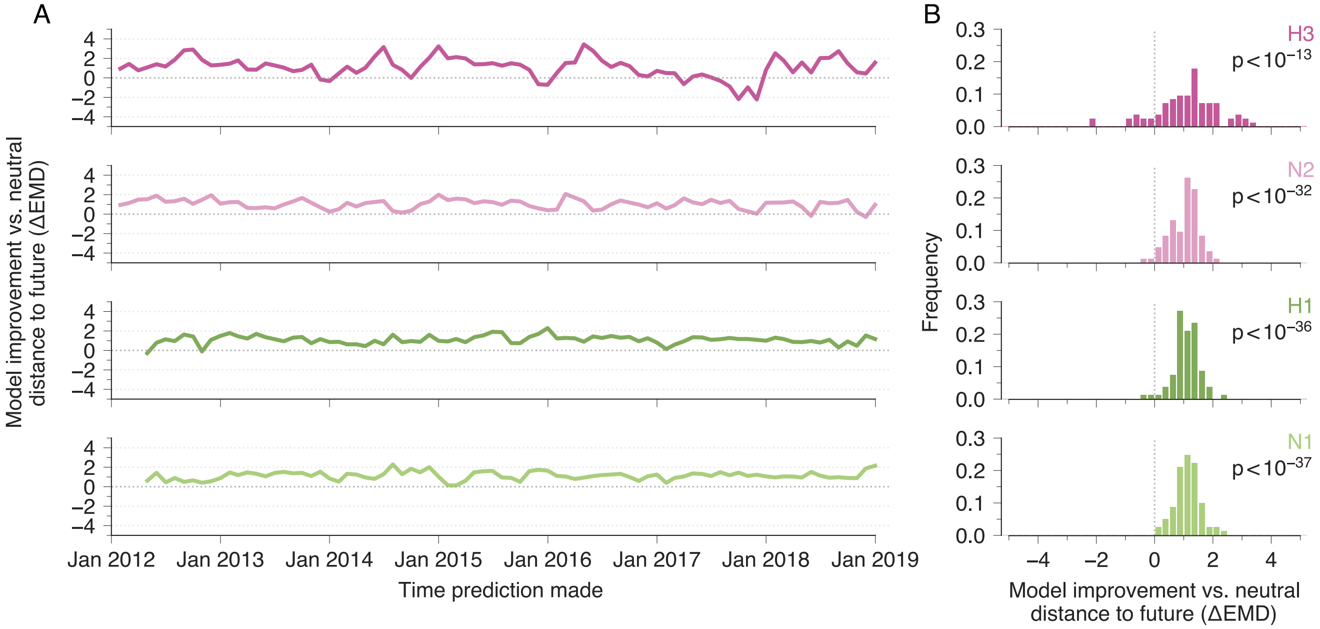


**Fig. S7. Simulated vs. neutral predictions of the composition of the IAV population 6 months in the future.** (**A**) Simulated predictions of the future composition of the IAV population outperform neutral ones, as measured by the difference in earth mover's distances ($\Delta\mathrm{EMD} = \mathrm{EMD}_{\mathrm{neutral}} - \mathrm{EMD}_{\mathrm{simulation}}$) between the true population and neutral vs. simulated ones. (**B**) Histogram of $\Delta\mathrm{EMD}$ values across time. For each IAV subtype and surface protein, our simulations improve significantly over neutral ones (Student's t-test).

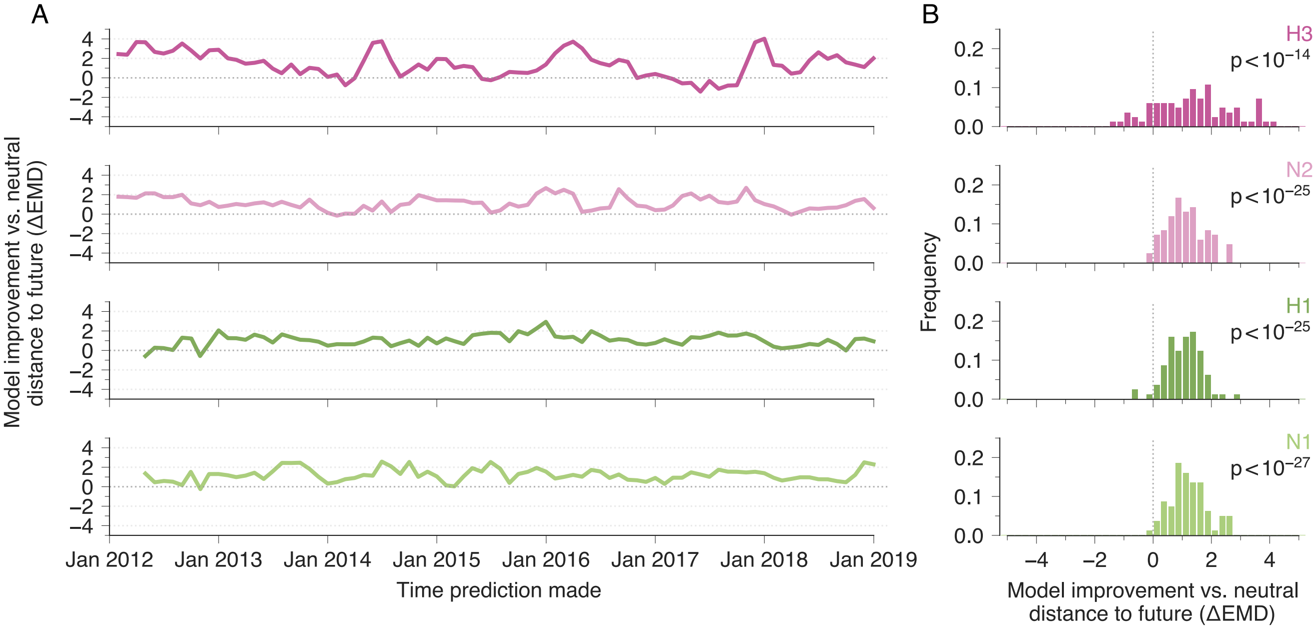


**Fig. S8. Simulated vs. neutral predictions of the composition of the IAV population 12 months in the future.** (**A**) Simulated predictions of the future composition of the IAV population outperform neutral ones, as measured by the difference in earth mover's distances ($\Delta\mathrm{EMD} = \mathrm{EMD}_{\mathrm{neutral}} - \mathrm{EMD}_{\mathrm{simulation}}$) between the true population and neutral vs. simulated ones. (**B**) Histogram of $\Delta\mathrm{EMD}$ values across time. For each IAV subtype and surface protein, our simulations improve significantly over neutral ones (Student's t-test).

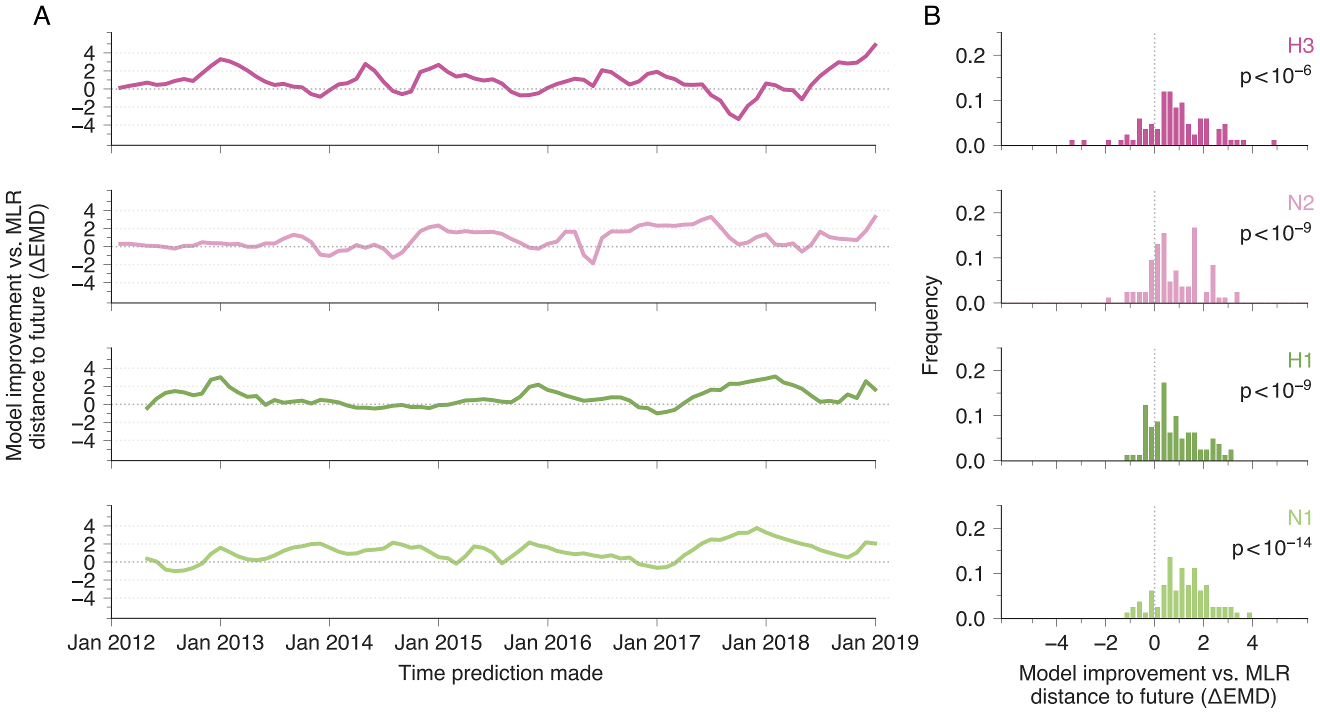


**Fig. S9. Simulated vs. MLR predictions of the composition of the IAV population 6 months in the future.** (**A**) Simulated predictions of the future composition of the IAV population outperform MLR ones, as measured by the difference in earth mover's distances ($\Delta\mathrm{EMD} = \mathrm{EMD}_{\mathrm{MLR}} - \mathrm{EMD}_{\mathrm{simulation}}$) between the true population and MLR vs. simulated ones. (**B**) Histogram of $\Delta\mathrm{EMD}$ values across time. For each IAV subtype and surface protein, our simulations improve significantly over MLR ones (Student's t-test).

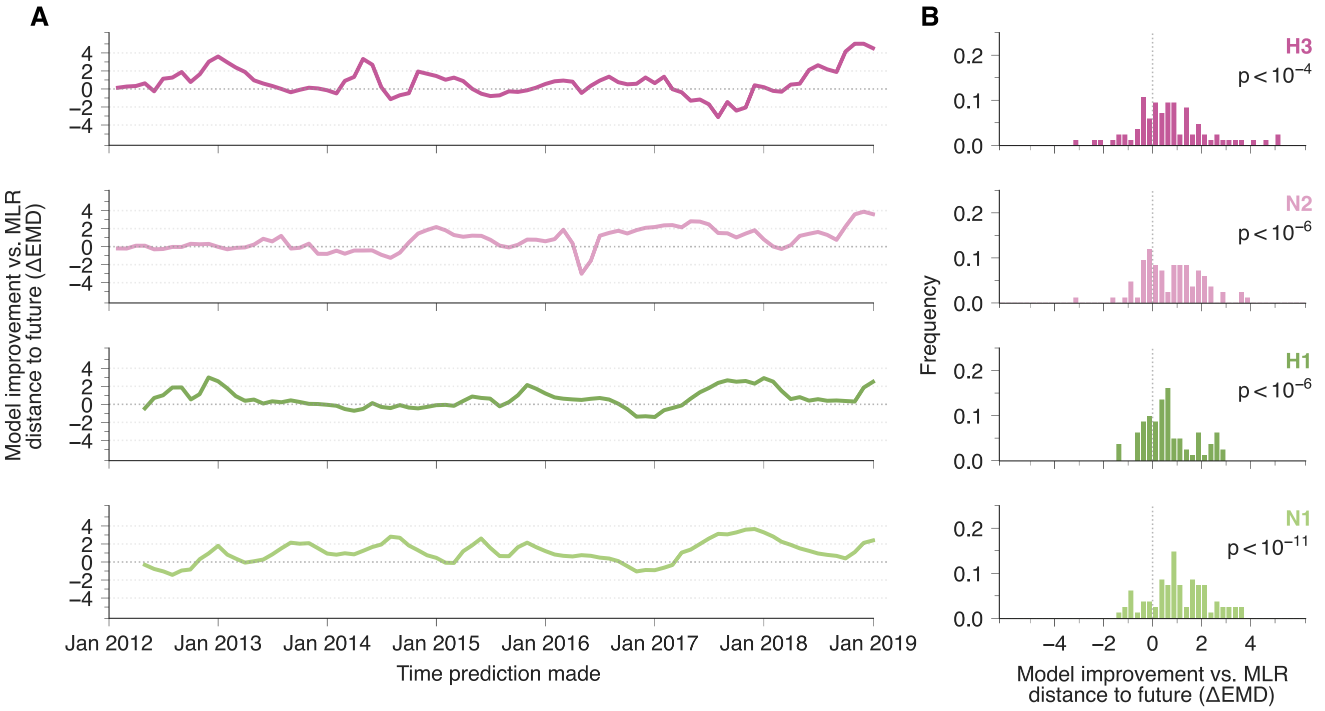


**Fig. S10. Simulated vs. MLR predictions of the composition of the IAV population 9 months in the future.** (**A**) Simulated predictions of the future composition of the IAV population outperform MLR ones, as measured by the difference in earth mover's distances ($\Delta\mathrm{EMD} = \mathrm{EMD}_{\mathrm{MLR}} - \mathrm{EMD}_{\mathrm{simulation}}$) between the true population and MLR vs. simulated ones. (**B**) Histogram of $\Delta\mathrm{EMD}$ values across time. For each IAV subtype and surface protein, our simulations improve significantly over MLR ones (Student's t-test).

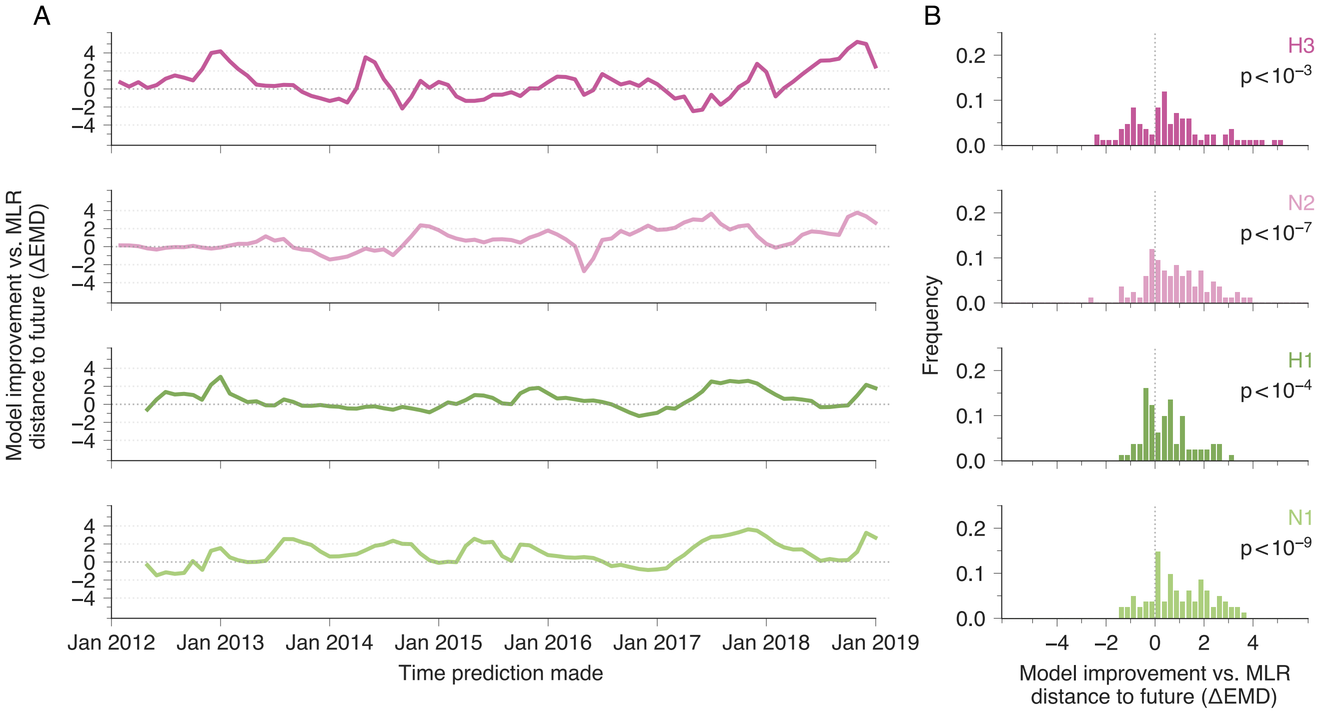


**Fig. S11. Simulated vs. MLR predictions of the composition of the IAV population 12 months in the future.** (**A**) Simulated predictions of the future composition of the IAV population outperform MLR ones, as measured by the difference in earth mover's distances ($\Delta\mathrm{EMD} = \mathrm{EMD}_{\mathrm{MLR}} - \mathrm{EMD}_{\mathrm{simulation}}$) between the true population and MLR vs. simulated ones. (**B**) Histogram of $\Delta\mathrm{EMD}$ values across time. For each IAV subtype and surface protein, our simulations improve significantly over MLR ones (Student's t-test).

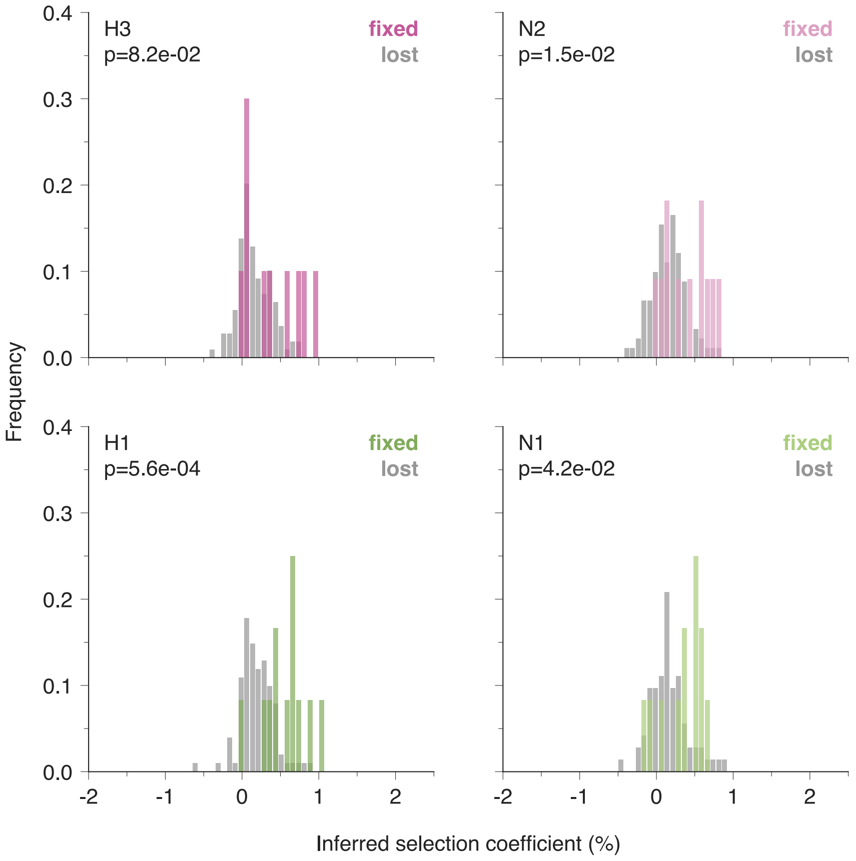


**Fig. S12. Mutations that fix tend to have larger inferred fitness contributions than mutations that are lost.** Distributions of inferred mutation-level fitness contributions, calculated using complete evolutionary trajectories, are shown separately for mutations that ultimately fixed and those that were lost. Fixed mutations generally have larger inferred fitness contributions, although the strength of this association varies among proteins (Student's t-test).

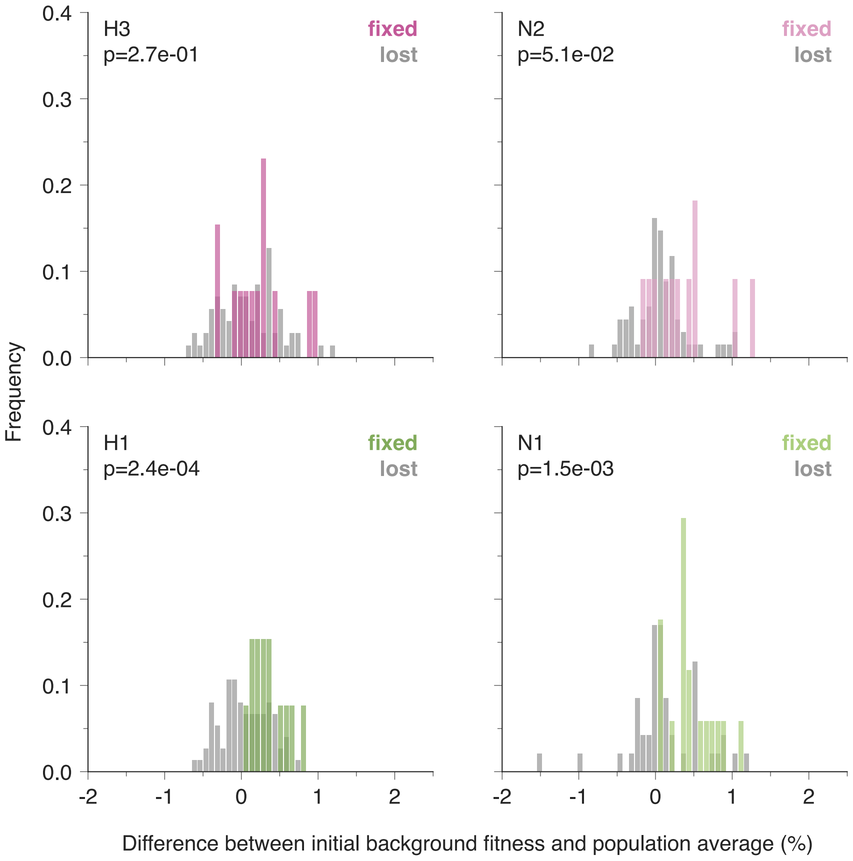


**Fig. S13. Mutations that fix preferentially arise on fitter sequence backgrounds.** The distributions of sequence background fitness, measured relative to the average fitness of the circulating population when each mutation first crossed the analysis threshold (**Methods**), are shown separately for mutations that ultimately fixed and those that were lost. Mutations that fixed generally occurred on fitter backgrounds.

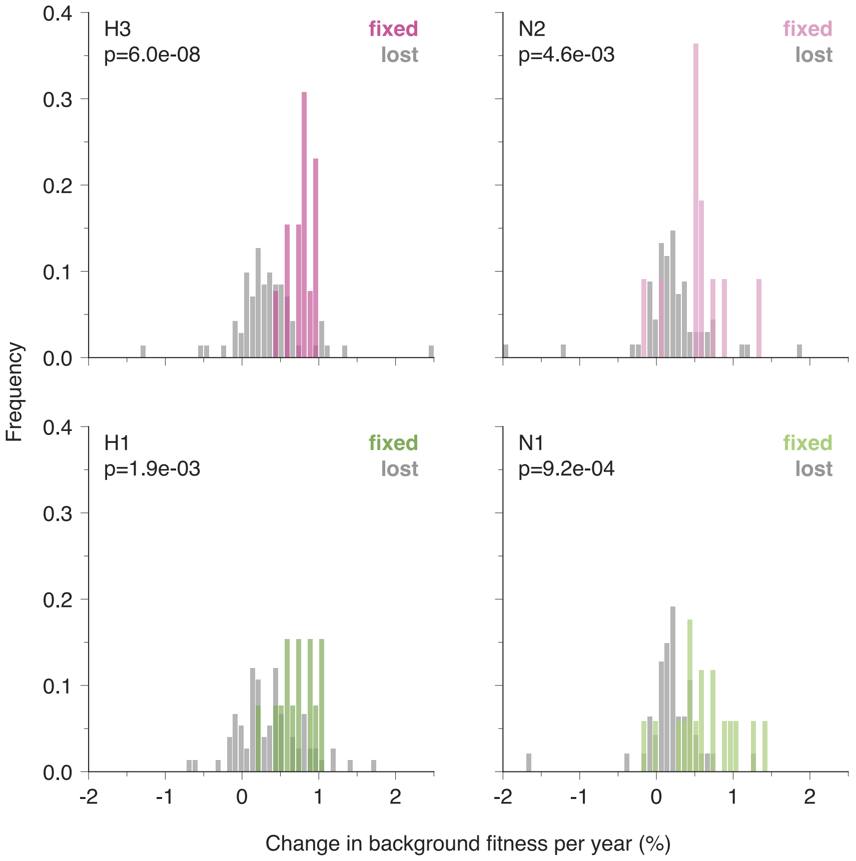


**Fig. S14. Sequence backgrounds carrying mutations that fix continue to increase in fitness more rapidly.** Distributions of the subsequent rates of change in sequence background fitness are shown separately for mutations that ultimately fixed and those that were lost. Backgrounds carrying mutations that fixed generally accumulated fitness more rapidly during subsequent evolution.

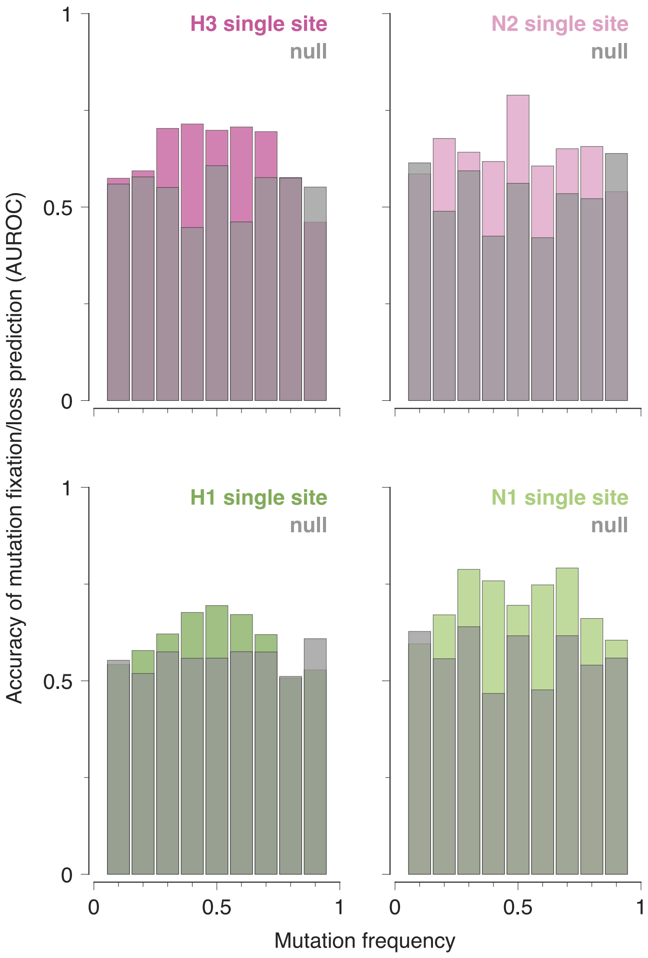


**Fig. S15. Genetic background improves predictions of mutation fate.** Accuracy of mutation fixation and loss predictions using the simplified single site model compared with the neutral model, measured by AUROC as a function of mutation frequency. Although the single site model improves upon neutral predictions, it remains less accurate than the full sequence-structured model shown in **Fig. S5**.

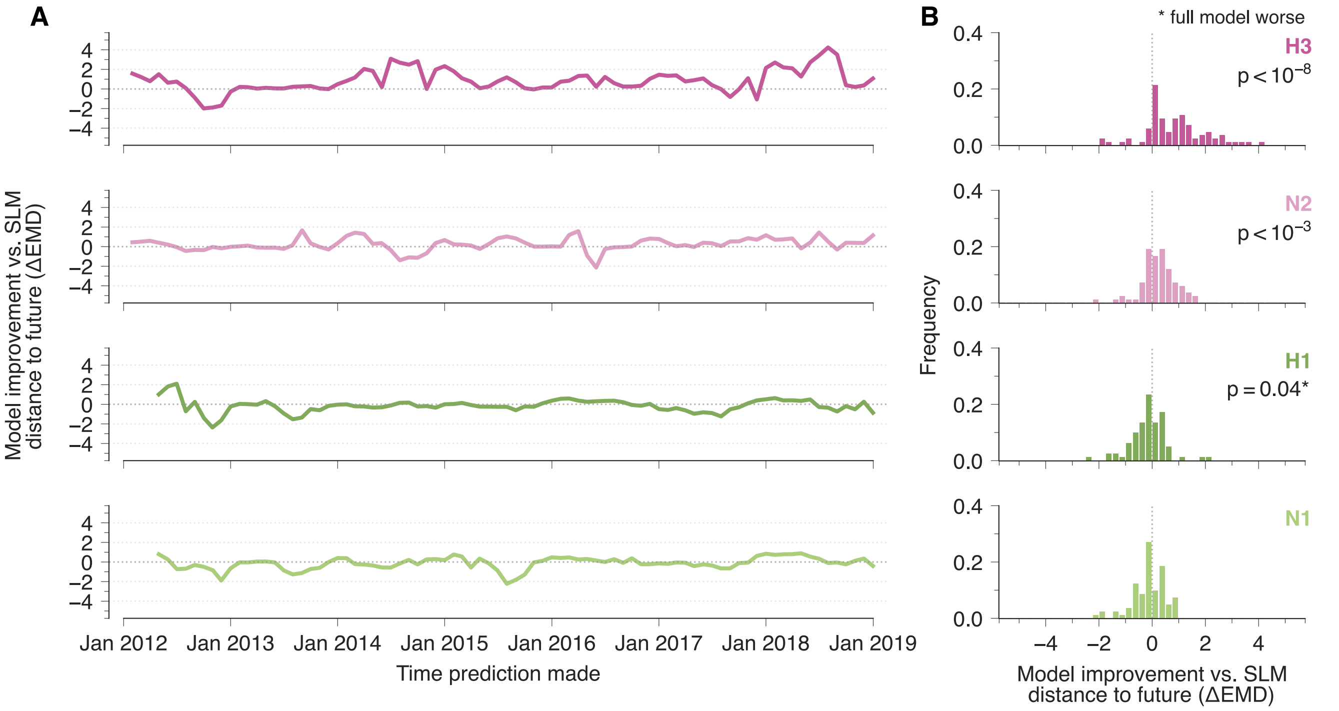


**Fig. S16. Simulated predictions of the composition of the IAV population 6 months in the future, comparing the full model versus a simplified, single site version.** (**A**) For H3N2, simulated predictions of the future composition of the IAV population using the full model typically outperform ones using the single site model, as measured by the difference in earth mover's distances ($\Delta\mathrm{EMD} = \mathrm{EMD}_{\text{single site}} - \mathrm{EMD}_{\text{full simulation}}$) between the true population and single site vs. full model simulated ones. (**B**) Histogram of $\Delta\mathrm{EMD}$ values across time. For both H3 and N2, the full model simulations improve significantly over single site ones (Student's t-test). For H1, the single site model actually performs slightly better than the full model.

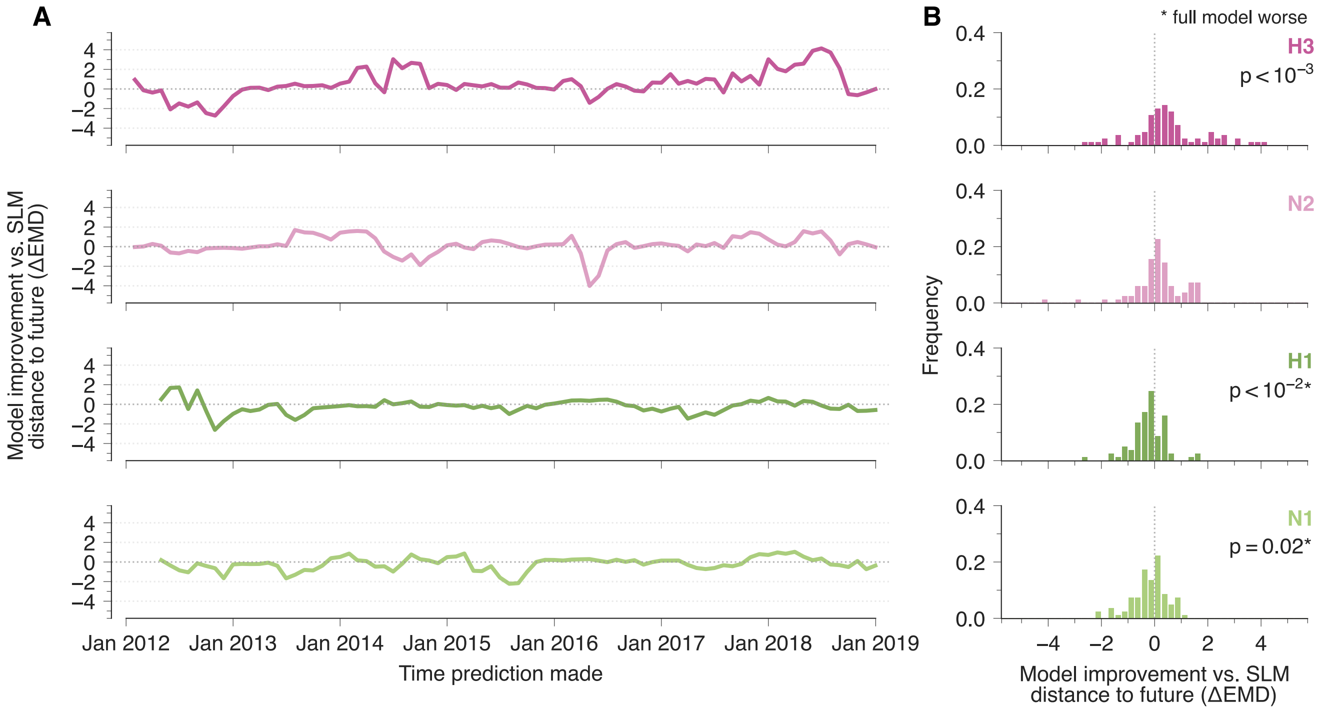


**Fig. S17. Simulated predictions of the composition of the IAV population 9 months in the future, comparing the full model versus a simplified, single site version.** (**A**) Comparative predictions of the future composition of the IAV population using the difference in earth mover's distances ($\Delta\mathrm{EMD} = \mathrm{EMD}_{\mathrm{single\ site}} - \mathrm{EMD}_{\mathrm{full\ simulation}}$) between the true population and single site vs. full model simulated ones. (**B**) Histogram of $\Delta\mathrm{EMD}$ values across time. For H3, the full model simulations improve significantly over single site ones (Student's t-test). For H1 and N1, the simpler single site model performs significantly better.

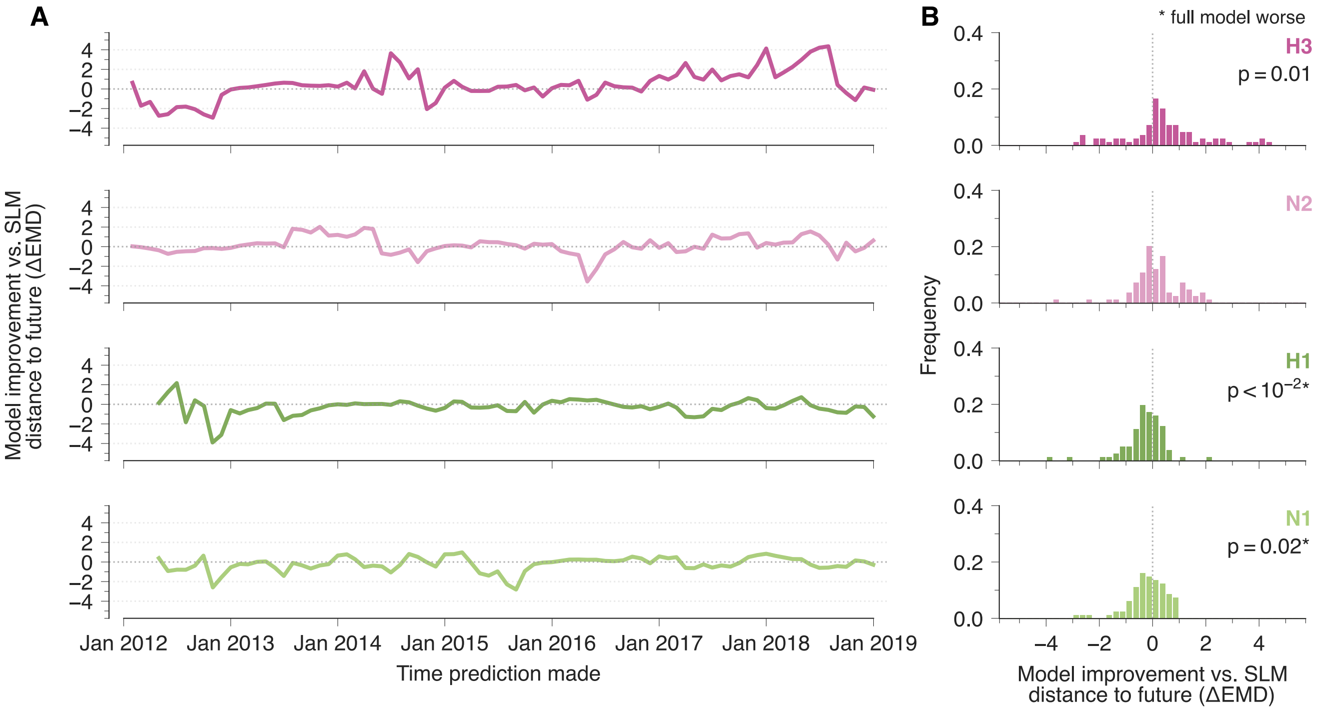


**Fig. S18. Simulated predictions of the composition of the IAV population 12 months in the future, comparing the full model versus a simplified, single site version.** (**A**) Simulated predictions of the future composition of the IAV population using the full model typically outperform ones using the single site model, as measured by the difference in earth mover's distances ($\Delta\mathrm{EMD} = \mathrm{EMD}_{\mathrm{single\ site}} - \mathrm{EMD}_{\mathrm{full\ simulation}}$) between the true population and single site vs. full model simulated ones. (**B**) Histogram of $\Delta\mathrm{EMD}$ values across time. As for the 9 month time horizon, the full model simulations improve significantly over single site ones for H3 (Student's t-test), while the signle site model performs better for H1 and N1. For N2, there is no significant difference in performance between the models.

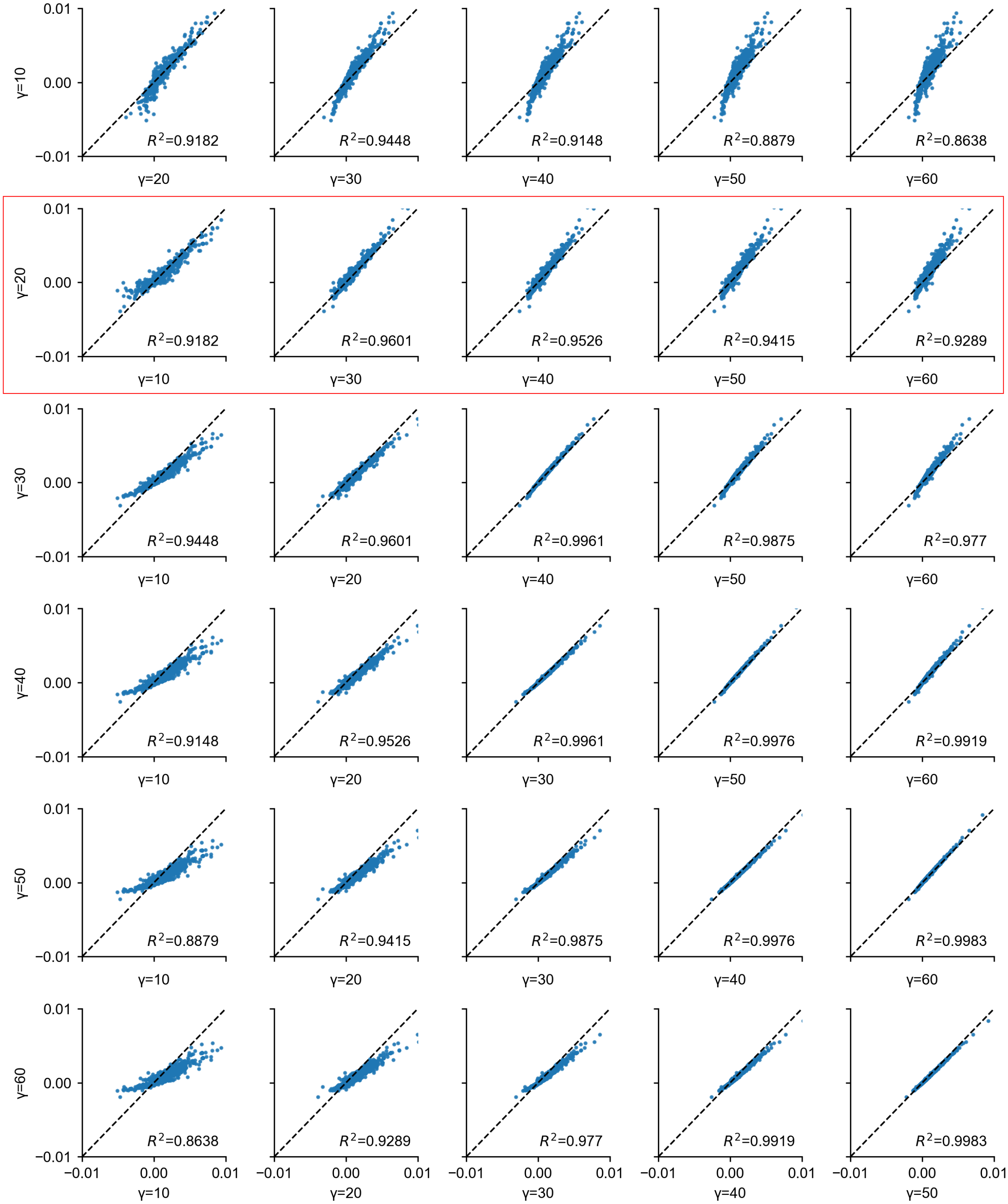


**Fig. S19. Inferred selection coefficients are robust to different values of the regularization parameter** $\gamma$**.** Pairwise comparisons of inferred selection coefficients for different regularization strengths. For small regularization ($\gamma = 10$), the inferred coefficients show greater variability, whereas for $\gamma \geq 20$ the estimates are highly correlated and exhibit consistent rank ordering. This indicates that the inference is stable with respect to the choice of $\gamma$, and that $\gamma = 20$ lies within the regime where results are largely insensitive to the precise regularization value.

| Domain | Mutation | Selection (%) | Structural Region | Antigenic/functional annotation | Literature support | Reported phenotype |
|---|---|---|---|---|---|---|
| HA1 | T131K | 1.36 | Head | Site A; RBS rim;serum escape | ESC-B; REG-C [48–51] | Diminished neutralization by Switzerland/2013 and Texas/2012-like sera within the 3C.2a2 T131K/R142K/R261Q haplotype; not tested alone. |
| HA2 | M346L | 1.02 | Stalk | – | – | No curated evidence. |
| HA1 | N189K | 1.00 | Head | Koel-7; site B; serum escape | POS-A; ESC-B; REG-C [49–53] | Koel/escape position; exact change untested. |
| HA1 | F159Y | 0.84 | Head | Koel-7; site B; serum escape; adjacent to N158 glycan | POS-A; ESC-B; REG-C [48,49,52–54] | Characterizes 3C.2a/3C.2a1 viruses; exact change untested. |
| HA1 | A198P | 0.74 | Head | Site B | REG-C [48,50,51] | Classical-site member; 3C.3a viruses carrying the substitution (as S198P). |
| HA2 | I478M | 0.74 | Stalk | – | – | No curated evidence. |
| HA1 | S45N | 0.72 | N-stalk | Site C; glycosylation motif gain | REG-C [50,51] | Classical-site member; exact change untested. |
| HA1 | N145S | 0.67 | Head | Koel-7; site A; serum/mAb escape | MUT-A; POS-A; ESC-B; REG-C [49–52,54,55] | Koel/escape position; Reverse-genetics N145S virus reduced serum inhibition. |
| HA1 | K158N | 0.57 | Head | Koel-7; site B; glycosylation motif gain | MUT-B; POS-A; REG-C [50–52,56] | Altered H3N2 antibody recognition. |
| HA2 | D489N | 0.56 | Stalk | Glycosylation motif gain | – | No curated evidence. |
| HA1 | F193S | 0.52 | Head | Koel-7; site B; serum escape | MUT-B; POS-A; ESC-B [48,49,52,53] | Reduced site-B mAb inhibition; repeatedly linked to antigenic distancing in natural 3C.3a, 3C.2a and 3C.2a1 viruses. |
| HA1 | N121K | 0.50 | Head | Site D | REG-C [50,51] | Exact change untested. |
| HA1 | L3I | 0.47 | N-stalk | – | – | No curated evidence. |
| HA1 | K92R | 0.46 | Head | F005-126 footprint; site E | FP-B; REG-C [48,50,51,57] | Antibody-contact position; exact change untested. |
| HA1 | T135K | 0.46 | Head | RBS; site A; N133 sequon loss; serum escape | POS-B; ESC-B; REG-C [48–51,58,59] | RBS/escape position; removes the N133 sequon; some T135K viruses showed reduced neutralisation by Hong Kong/2014-like antiserum. |
| HA1 | T128A | 0.45 | Head | Site B | REG-C [50,51] | Exact change untested. |
| HA1 | F159S | 0.43 | Head | Koel-7; site B; serum escape | MUT-B; POS-A; ESC-B [49,52–54] | F159S reduced antisera and mAb recognition. |
| HA1 | N171K | 0.43 | Head | F005-126 footprint; site D | FP-B; REG-C [50,51,57] | Antibody-contact position; exact change untested. |
| HA1 | K2E | 0.41 | N-stalk | – | – | No curated evidence. |
| HA1 | N31S | 0.39 | N-stalk | – | – | No curated evidence. |
| HA1 | N278K | 0.39 | C-stalk | Site C | REG-C [50,51] | Exact change untested. |

**Table 1. Top 20 positively selected mutations in H3N2 HA, ranked by inferred selection coefficient.** Positions use mature H3 HA numbering, excluding the 16-amino-acid signal peptide, anchored to A/Aichi/2/68. Evidence codes are: **MUT**, the same amino acid substitution was experimentally studied; **POS**, the residue was experimentally implicated but not necessarily for the listed substitution; **FP**, a structurally resolved antibody footprint; **ESC**, antibody- or serum-escape evidence, including mutational antigenic profiling; and **REG**, membership in an established antigenic or functional region. Grades modify the preceding code: **A**, direct experimental evidence at the mapped H3N2 residue; **B**, evidence from another strain or sequence background, or context-dependent structural or high-throughput evidence; and **C**, region membership without substitution-specific validation. Position or region membership does not establish an effect for the exact substitution.

| Domain | Mutation | Selection (%) | Structural Region | Antigenic/functional annotation | Literature support | Reported phenotype |
|---|---|---|---|---|---|---|
| HA1 | S164T | 1.00 | Head | Immunodominant site Sa; N162 glycosylation module | POS-B; REG-C [60,61] | Converts the N162 sequon from NQS to NQT and was proposed to stabilise N162 glycosylation; antigenic effect untested. |
| HA1 | S185I | 0.94 | Head | RBS; sites Sb | POS-A; REG-C [61–63] | Defining substitution of the 6B.1A5A lineage, together with N129D and N260D |
| HA1 | K163Q | 0.86 | Head | Site Sa; lateral patch, N162 glycosylation module | MUT-A; ESC-A; REG-C [60,61,64,65] | Reduced antibody recognition in H1N1; permissive for complete N162 glycosylation. |
| HA1 | I324V | 0.85 | C-stalk | – | – | No curated evidence. |
| HA1 | N129D | 0.82 | Head | OREO epitope (VR1); Sa-proximal | MUT-A; ESC-A; POS-A; REG-C [66–68] | Selected as a human-mAb escape mutant and increased $\alpha$2,6 receptor binding without loss of replication in human cells. |
| HA1 | S74R | 0.77 | Head | site Cb; escape position | POS-B; ESC-B; REG-C [61] | S74G escaped in PR8; exact change untested. |
| HA2 | S451T | 0.76 | Stalk | T-cell peptide RSQLKNNAK | REG-C [69] | T-cell peptide member; exact change untested. |
| HA1 | Q189E | 0.73 | Head | RBS; sites Sb | POS-A; ESC-B; REG-C [61–63] | Other changes escaped antibodies; observed in 6B.1A5A descendants; exact change untested. |
| HA1 | K283E | 0.69 | C-stalk | – | – | No curated evidence. |
| HA2 | E499K | 0.68 | Stalk | – | – | No curated evidence. |
| HA2 | E374K | 0.66 | Stalk | Pa-associated combination | MUT-B [62] | Reduced recognition only as N125D+E374K combination. |
| HA1 | S162N | 0.66 | Head | site Sa; lateral patch; glycosylation motif gain | MUT-A; ESC-A; REG-C [60,61,65] | Reduced binding of lateral-patch antibodies; creates the N162 sequon. |
| HA2 | V520A | 0.62 | Membrane | – | – | No curated evidence. |
| HA2 | S451N | 0.61 | Stalk | T-cell peptide RSQLKNNAK | REG-C [69] | T-cell peptide member; exact change untested. |
| HA1 | D187A | 0.58 | Head | site Sb; escape position | POS-A; ESC-A; REG-C [61,70] | D187E escaped human mAb 5E4; observed in 6B.1A5A descendants; exact change untested. |
| HA1 | T120A | 0.57 | Head | – | – | No curated evidence. |
| HA1 | I216K | 0.57 | Head | N162 glycan occupancy determinant | POS-B [60] | Occupancy of the N162 sequon requires T216; the derived K (arising as T216K) is predicted to reduce occupancy but is untested. |
| HA2 | I404L | 0.57 | Stalk | – | – | No curated evidence. |
| HA1 | S183P | 0.54 | Head | RBS; site Ca2; Sb escape position | MUT-A; POS-A; REG-C [62,63,71,72] | Increased binding to several $\alpha$2,6-linked glycans. |
| HA1 | K130N | 0.51 | Head | Site Pa; OREO epitope (VR1) | POS-A; REG-C [62,68] | Mapped antigenic/OREO position; exact change untested. |

**Table 2. Top 20 positively selected mutations in A/H1N1 HA, ranked by inferred selection coefficient.** Positions use mature H1 numbering, excluding the 17-amino-acid signal peptide, anchored to A/California/07/2009. Evidence codes and grades follow Table 1. A dash indicates no matching curated literature record, and position- or region-level evidence does not establish an effect for the exact substitution.

| Domain | Mutation | Selection (%) | Structural Region | Antigenic/functional annotation | Literature support | Reported phenotype |
|---|---|---|---|---|---|---|
| NA | E344K | 0.83 | Head | E344 determinant; charge-balanced region; DA03E17 footprint | MUT-A; POS-A; FP-B; REG-C [73–77] | Reduced antisera recognition in contemporary H3N2. |
| NA | T267K | 0.80 | Head | Link to the N245 glycan haplotype | MUT-B [78] | Tested only within an S245N/S247T/T267K construct in A/Hong Kong/4801/2014 N2, where inhibition by active-site antibodies was altered; the contribution of T267K was not resolved from the N245 glycan. |
| NA | I469T | 0.79 | Head | – | – | No curated evidence. |
| NA | S367N | 0.74 | Head | 2SBS 370 loop; potential N367 sequon with K369T | MUT-B; POS-A; REG-C [74,76,77,79] | Disrupted 2SBS-dependent binding and cleavage of multivalent receptors in a reconstructed 1957 human N2, with the reciprocal N367S restoring both; also reduced hemadsorption and altered catalytic activity. Not tested in a contemporary seasonal N2. |
| NA | I57M | 0.72 | Stalk | – | – | No curated evidence. |
| NA | L338S | 0.70 | Head | Lateral antigenic loop | REG-C [76] | Antigenic-loop member; exact change untested. |
| NA | K369T | 0.67 | Head | 2SBS 370 loop; potential N367 sequon with S367N | POS-A; REG-C [74,76,77,80] | Same 2SBS loop as residue 367; individual effects on receptor processing not isolated. |
| NA | E221D | 0.61 | Head | Mem5 footprint; 220-loop | POS-B; FP-B; REG-C [76,81,82] | Other residue backgrounds mediate Mem5 escape; exact change untested. |
| NA | P468H | 0.59 | Head | Post-2016 antigenic-drift determinant | MUT-A; POS-A [78] | Contributed to post-2016 N2 antigenic drift. |
| NA | N402D | 0.59 | Head | 2SBS 400 loop; Colman segment IV | REG-C [77,79] | Historical antigenic-region member; Located in the 400 loop of the 2SBS; functional and antigenic effects untested. |
| NA | I176M | 0.53 | Head | – | – | No curated evidence. |
| NA | I26V | 0.52 | Transmembrane | – | – | No curated evidence. |
| NA | N161S | 0.49 | Head | – | – | No curated evidence. |
| NA | V303I | 0.45 | Head | – | – | No curated evidence. |
| NA | S247T | 0.45 | Head | N245 glycan determinant; DA03E17 footprint | MUT-B; POS-A; FP-B [75,78] | Promoted N245 glycosylation with S245N. |
| NA | P386S | 0.40 | Head | – | – | No curated evidence. |
| NA | I392T | 0.38 | Head | – | – | No curated evidence. |
| NA | I469V | 0.36 | Head | – | – | No curated evidence. |
| NA | N329S | 0.35 | Head | N329 glycosylation determinant (sequon loss); lateral antigenic loop; Colman segment I | POS-A; REG-C [73,77,81] | Removes the N329 sequon; experimental manipulation of glycosylation at 329 altered inhibition by human and ferret antisera, exact change untested. |
| NA | L81P | 0.34 | Head | – | – | No curated evidence. |

**Table 3. Top 20 positively selected mutations in H3N2 N2, ranked by inferred selection coefficient.** Positions use full-length N2 numbering. Literature annotations are anchored to A/Memphis/31/98 and mapped to the A/Indiana/10/2011 sequence alignment. Evidence codes and grades follow Table 1; **OBS** denotes observational or evolutionary association without direct substitution-level validation. A dash indicates no matching curated literature record, and position- or region-level evidence does not establish an effect for the exact substitution.

| Domain | Mutation | Selection (%) | Structural Region | Antigenic/functional annotation | Literature support | Reported phenotype |
|---|---|---|---|---|---|---|
| N1 | V13I | 0.81 | Transmembrane | Fixed post-2009 substitution | OBS [83] | No curated evidence. |
| N1 | I389K | 0.73 | Head | CD6 inter-protomer footprint; historical 387–395 antigenic region | FP-B ; REG-C [84,85] | CD6-contact position; exact change untested. |
| N1 | N386K | 0.70 | Head | $Ca^{2+}$-binding-site residue; potential sequon loss; post-2009 drift | MUT-A; POS-A; OBS [83,86] | Increased activity against multivalent substrates; removes a potential sequon. |
| N1 | K432E | 0.63 | Head | 2SBS 430 loop;Antigenicity classifier; DA03E17 footprint | MUT-A; POS-A; FP-B; OBS [75,83,86–88] | Largest single-step antigenic change in a chronological reconstruction of A/H1N1 N1; also altered enzymatic properties. |
| N1 | N369K | 0.61 | Head | 2SBS 370 loop; HF5 escape position | MUT-A; ESC-A; OBS [83,85,86] | Reduced HF5 inhibition in a CA/09 background; increased cleavage of fetuin, altered calcium-dependent thermostability, and produced a relatively large single-step antigenic change. |
| N1 | I106V | 0.61 | Head | Post-2009 drift-associated position | OBS [86] | Drift-associated sequence change; phenotype untested. |
| N1 | V81A | 0.60 | Stalk | – | – | No curated evidence. |
| N1 | N449D | 0.59 | Head | CD6 footprint; N449D/D451G escape pair | MUT-A; FP-B [85] | Limited alone; with D451G abolished CD6 binding. |
| N1 | G77R | 0.54 | Stalk | – | – | No curated evidence. |
| N1 | N270K | 0.53 | Head | CD6 footprint; human-mAb escape position | FP-B; ESC-B; OBS [83,85,86,89] | Tested only as the V264I/N270K pair, which increased cleavage of $\alpha$2,6-linked sialic acids; escape at this position was mediated by other changes. |
| N1 | V241I | 0.51 | Head | Post-2009 drift-associated position | OBS [86] | Drift-associated sequence change; phenotype untested. |
| N1 | Y66F | 0.51 | Stalk | – | – | No curated evidence. |
| N1 | I117M | 0.50 | Head | – | – | No curated evidence. |
| N1 | N397K | 0.50 | Head | Historical group-E epitope | POS-A; REG-C [84] | Experimentally mapped epitope position; exact change untested. |
| N1 | M19T | 0.49 | Transmembrane | – | – | No curated evidence. |
| N1 | N222K | 0.44 | Head | – | – | No curated evidence. |
| N1 | S52N | 0.44 | Stalk | Glycosylation motif gain | – | No curated evidence. |
| N1 | I467V | 0.43 | Head | – | – | No curated evidence. |
| N1 | F74S | 0.40 | Stalk | Glycosylation motif gain | – | No curated evidence. |
| N1 | N200S | 0.40 | Head | DA03E17 footprint | OBS; FP-B [75,86] | Drift-associated active-site-adjacent change; causality untested. |

**Table 4. Top 20 positively selected mutations in A/H1N1 N1 protein, ranked by inferred selection coefficient.** Positions use full-length N1 numbering anchored to A/California/07/2009. Literature coordinates reported in conventional NA numbering were mapped to the same alignment. Evidence codes and grades follow Table 1; **OBS** denotes an observational or evolutionary association without direct substitution-level validation. A dash indicates no matching curated literature record, and position- or region-level evidence does not establish an effect for the exact substitution.
